# SARS-CoV-2 saltational events are recurrent and trace to persistent human infections

**DOI:** 10.64898/2026.06.18.733214

**Authors:** Cécile Tran-Kiem, Kathryn Kistler, Ryan Hisner, Trevor Bedford

## Abstract

SARS-CoV-2 evolution is characterized by gradual mutation accumulation but has been punctuated by rare yet impactful highly mutated variants. Whether such saltational jumps are a broad feature of SARS-CoV-2 evolution or rare anomalies remains unclear. We systematically investigate SARS-CoV-2 saltational evolution by developing a scalable framework to detect saltational events from 4.4 million high-quality viral genomes. Saltational events occurred at low but detectable rates during the pandemic and post-pandemic periods and across geographies. Their mutational signature closely matches that seen in persistent human infections but is inconsistent with the signatures of mink or deer infections. This points to persistent infection, rather than reverse zoonosis, as their primary source. While most saltational events lack evidence of onward transmission, those that do tend to carry mutations found in successful clades. Our work demonstrates that the emergence of highly mutated SARS-CoV-2 variants reflects a recurrent evolutionary process, with implications for preparedness.

## Introduction

The accumulation of mutations in SARS-CoV-2 genomes typically occurs incrementally as the virus spreads throughout the population, with one substitution arising roughly every two weeks. However, SARS-CoV-2 evolution has been punctuated by the emergence of variants carrying unusually high numbers of mutations^1–4^, some of which have rapidly swept worldwide and generated “pandemics within the pandemic”^5^. Because of their major epidemiological consequences, these successful saltational events have attracted considerable attention.

Yet our understanding of SARS-CoV-2 saltational evolution remains largely based on a handful of high-profile variants of concern (VOCs), leaving it unclear whether such events were rare anomalies or a broader underappreciated feature of SARS-CoV-2 evolution. The unprecedented scale of genomic surveillance implemented in response to the pandemic offers an opportunity to address this question by quantifying how frequently saltational events occurred, how they were distributed across space and time and whether there are features associated with their successful spread. Such data may also provide insights into the mechanisms that gave rise to these events. Accelerated evolution during a subset of persistent SARS-CoV-2 infections has emerged as a very plausible hypothesis^6^, motivated by the overlap between mutations observed in chronically infected individuals and those found in VOCs^7^, with alternative explanations such as accumulation of mutation within animal reservoirs having little evidence to support them^7^.

Here, we perform a systematic analysis of SARS-CoV-2 saltational evolution from a large publicly available high-quality SARS-CoV-2 sequence dataset and phylogeny comprising 4.4 million samples^8^. We develop a scalable framework to detect saltational events, defined as branches in the phylogeny with strong evidence for positive selection and a high number of nonsynonymous mutations, and show they are not isolated anomalies but occur repeatedly during SARS-CoV-2 circulation. The criterion we use to define saltational branches does not incorporate temporal information. We find that such events have a distinct mutational signature that strongly resembles that observed in persistent SARS-CoV-2 infections. Finally, we investigate the extent to which saltational events seed onward transmission and identify mutational features associated with their successful spread.

## Results

### A scalable framework to detect saltational evolution in a mutation-annotated tree

To systematically study SARS-CoV-2 saltational evolution, we analyze a large SARS-CoV-2 phylogeny built from 4.4 million high-quality consensus sequences^8^, yielding a mutation-annotated tree with 5.3 million branches. We classify saltational branches as branches displaying both (i) an unusually high number of nonsynonymous mutations, (defined as branches with at least 5 nonsynonymous mutations, corresponding to greater than the 99.5th percentile of the distribution across all branches of the phylogeny, see Figure S1), and (ii) evidence of positive selection (Figure 1A). Because our definition of saltational branches only relies on substitution information and does not incorporate temporal information, identifying them only requires a mutation-annotated tree rather than a dated phylogeny. We infer positive selection at the gene or whole-genome level from an excess of nonsynonymous relative to synonymous mutations on a branch, accounting for nonsynonymous and synonymous mutation opportunities (*d*_*N*_*/d*_*S*_ ratio above 1). To do so, we develop a Bayesian framework to estimate branch- and gene-specific *d*_*N*_*/d*_*S*_ while accounting for sparse mutation counts. This approach allows us to account for uncertainty in branch-level *d*_*N*_*/d*_*S*_ estimates and to handle the fact that most branches contain few, if any, synonymous mutations. We then use Bayesian hypothesis testing to identify branches with strong evidence that *d*_*N*_*/d*_*S*_ > 1 at the gene or whole-genome level (see Methods).

**Figure 1:**
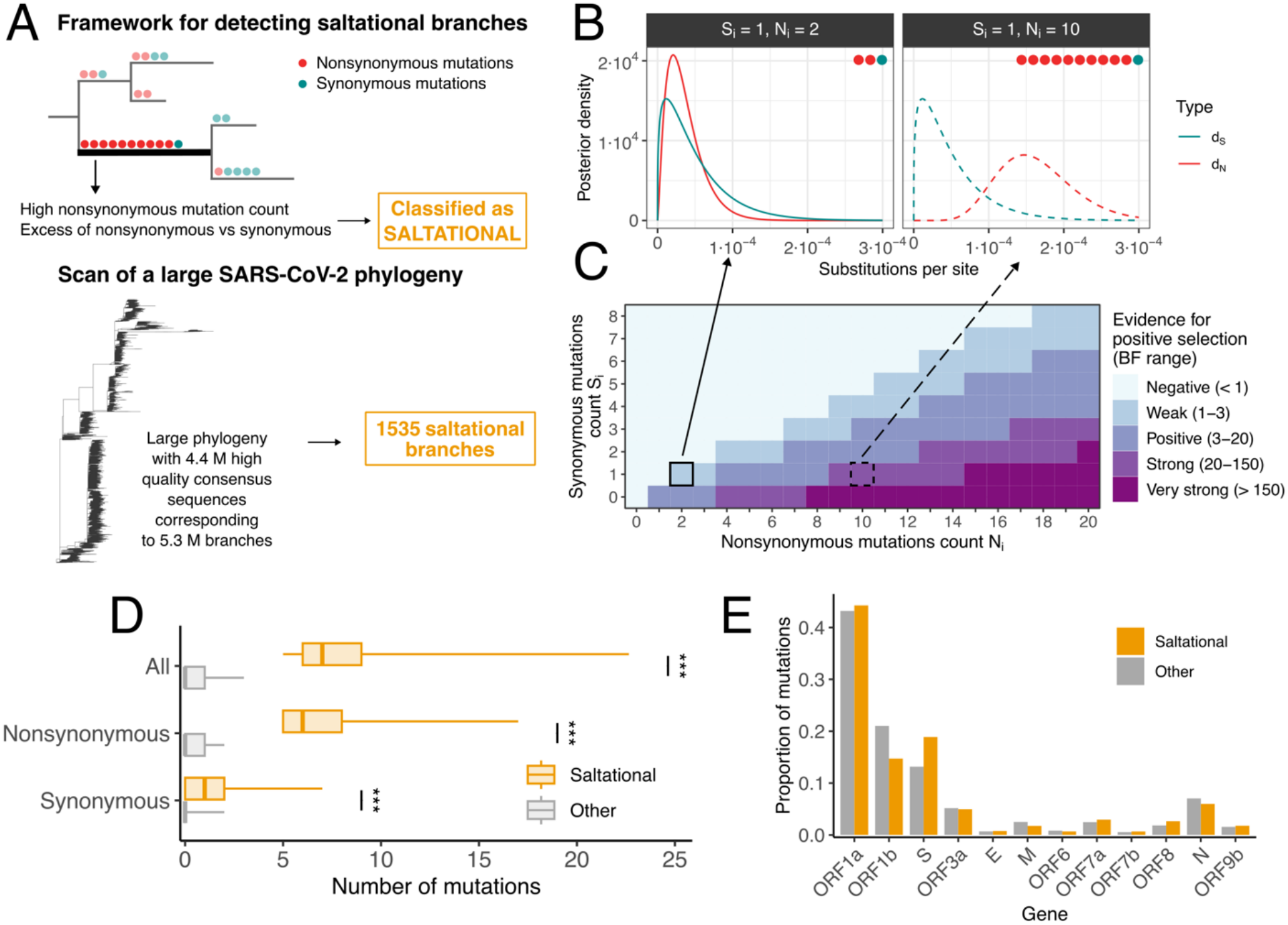
Framework for detecting saltational branches in a large SARS-CoV-2 phylogeny. **A**. Schematic overview of the framework used to classify branches as saltational, which we apply on a large SARS-CoV-2 phylogeny. **B**. Posterior distribution of nonsynonymous (*d*_*N*_) and synonymous (*d*_*S*_) divergence measured as substitutions per site on the spike (S) gene for a branch harboring 1 synonymous mutation and 2 or 10 nonsynonymous mutations. **C**. Strength of evidence for positive selection on the spike gene as a function of the number of nonsynonymous and synonymous mutations in spike occurring on a branch. The range in parenthesis in the legend indicates the range of the Bayes Factor for *d*_*N*_*/d*_*S*_ > 1 corresponding to each evidence category^9^. **D**. Number of mutations (total, nonsynonymous and synonymous) occurring on saltational compared with non-saltational branches. **E**. Proportion of mutations occurring on each gene on saltational and non-saltational branches. In D, boxplots indicate the 2.5th, 25th, 50th, 75th and 97.5th percentiles.

Figure 1B-C illustrates the model’s behavior. On a branch carrying 1 synonymous and 2 nonsynonymous mutations in the spike gene (S), we infer a posterior median of 3.7⋅10^-5^ synonymous mutations per site and a posterior median of 3.1⋅10^-5^ nonsynonymous mutations per site, resulting in little evidence for positive selection with d_N_/d_S_ estimated at 0.84 [95% credible interval: 0.086–14] and a Bayes Factor of 1.1. By contrast, on a branch carrying 10 nonsynonymous mutations in S, the posterior median of *d*_*N*_ increases to 1.6⋅10^-4^, resulting in strong support for positive selection with d_N_/d_S_ estimated at 4.2 [95% credible interval: 0.83–63] and a Bayes Factor of 31. Applying this framework to a tree built from 4.4 million samples collected between January 2020 and June 2024, we identify 1535 saltational branches. Compared with non-saltational branches, these branches carry substantially more mutations, especially nonsynonymous ones (Figure 1D), and are enriched in mutations in Spike (Figure 1E). Considering stricter thresholds for the definition of saltational evolution results in identifying fewer saltational branches (Figure S2). The number of branches classified as saltational is comparable to that obtained under a null model preserving substitution counts per branch where substitution types (synonymous or nonsynonymous) are redrawn based on the tree-wide gene-specific d_N_/d_S_ ratios (Figure S3). However, observed saltational branches carry more nonsynonymous and Spike mutations than saltational branches obtained from the null simulations.

Branches carrying a large number of nonsynonymous mutations could represent both a true biological signal or sequencing and bioinformatic errors. Relying on the Viridian tree, which was constructed to correct for known systematic assembly and bioinformatic errors, should improve our ability to identify a true biological signal^8^. To assess saltational branches quality, we run Nextclade on terminal saltational branches^10^ (Figure S4). As expected, saltational branches display a high number of private mutations. A few saltational branches were flagged as having many missing data. Overall, almost all terminal saltational branches showed no evidence of major sequence quality problems based on frameshifts, stop codons, mutation clusters and mixed sites. As a conservative sensitivity analysis, we reapply our saltational evolution criteria when removing mutations identified by Nextclade as occurring within mutation clusters, frameshifts regions, at mixed sites or downstream of stop codons. Of the 1,072 terminal saltational branches we identified, 1000 are still classified as saltational after this filtering. This suggests that the branches we identify are unlikely to be predominantly driven by poor sequence quality.

### Saltational evolution occurred at a low but detectable rate since SARS-CoV-2 emergence

We next investigate how the saltational branches we identify are distributed through space and time. We detect saltational branches in every year of the study period, on both terminal and internal branches of the phylogeny (Figure 2A). Their absolute count varies across years, with more events detected during periods where more sequences are collected. Overall, 0.03% of all branches are classified as saltational. Year-specific proportions are higher in 2023 and 2024 than in previous years (Figure 2B): whereas we identify 0.019% of branches as saltational in 2022 [95% CI: 0.017–0.021%], this proportion rises by ten-fold to 0.19% [95% CI: 0.14–0.25%] in 2024. This increase is also observed when subsampling the tree to the same number of samples per year (Figure S5), indicating that this trend cannot be only attributed to the lower number of sequences available in more recent years.

**Figure 2:**
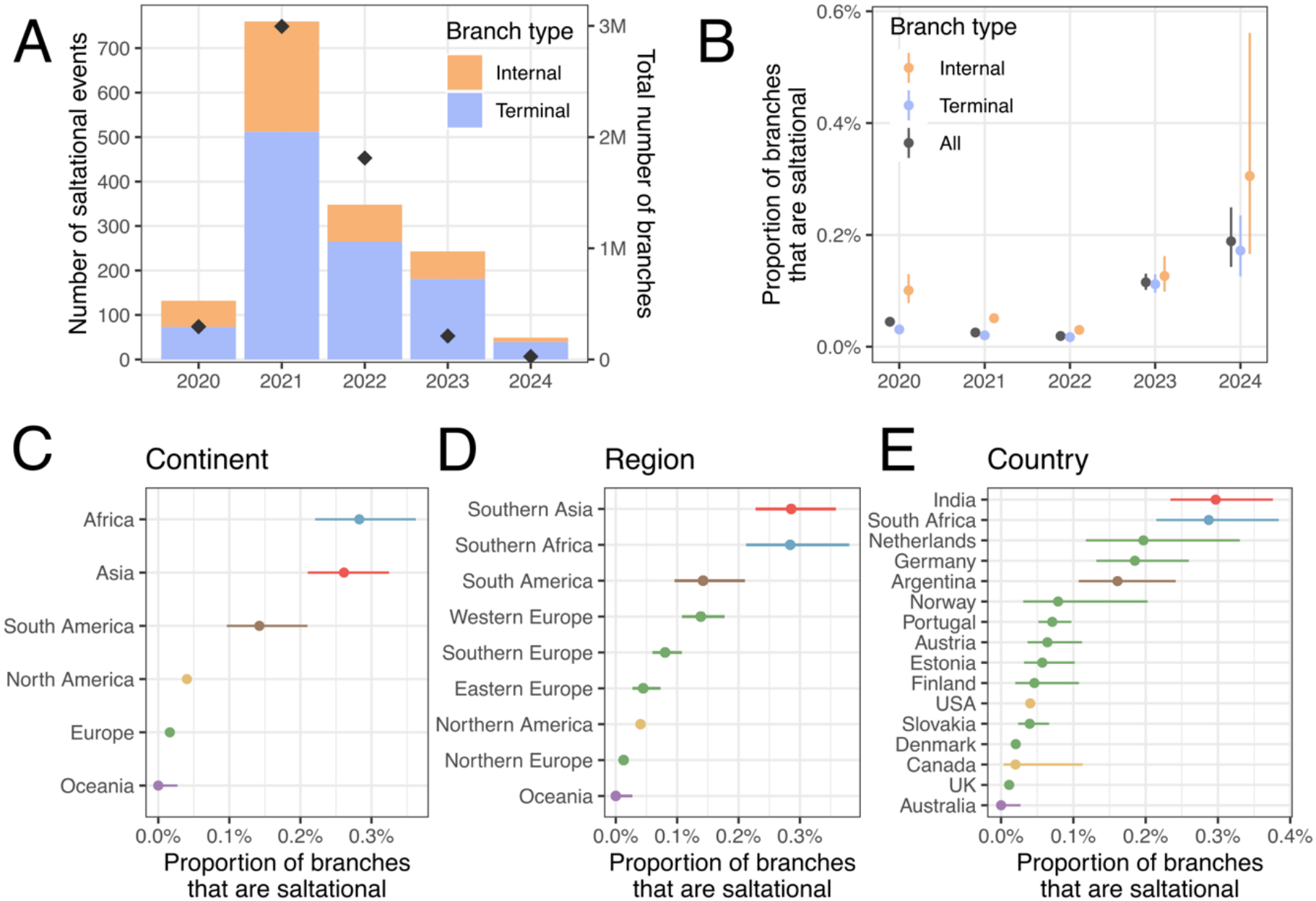
Spatiotemporal distribution of saltational SARS-CoV-2 branches. **A**. Number of saltational branches identified by year and branch type (bars with scale on the left). Diamonds depict the total number of branches for each year (right axis). **B**. Proportion of branches classified as saltational by year and branch type. Proportion of branches classified as saltational by **C**. continent, **D**. region and **E**. country. In B-E, segments indicate 95% Wilson confidence intervals.

We identify saltational branches across all continents represented in the dataset we studied, except Oceania (Figure 2C), highlighting their broad geographic distribution. Importantly, 99.6% of Oceanian sequences included in the tree were collected in 2020, a period during which both Australia and New Zealand (which represent almost all the Oceanian sequences in the analyses) implemented a zero-Covid strategy, leading to a very low infection burden. The lack of saltational events detected in this region thus likely reflects the limited opportunities for their emergence in the early stage of the pandemic when transmission was strongly suppressed. Looking at absolute counts, we detect the largest number of saltational events in the USA (784) and the UK (306), the two countries contributing the largest number of sequences in the studied dataset. The fact that we detect more saltational events in locations with higher sequencing (Figure S6) suggests that saltation is a general feature of SARS-CoV-2 evolution that can be observed when genomic surveillance is sufficiently intense. Focusing on regions with more than 5000 branches in the global phylogeny (see Methods for ancestral reconstruction), we find some variation in the proportion of branches that are saltational (Figure 2D-E), ranging from 0.0% in Australia [95% CI: 0.00–0.03%] to 0.3% in both India and South Africa [95% CI: 0.2–0.4% for South Africa and 0.2– 0.3% for India]. These observed differences could stem from heterogeneity in emergence opportunities and geographic variation in the detectability of saltational events. Taken together, these findings show that saltational evolution is a recurring feature of the COVID-19 pandemic with a broad geographic footprint.

### SARS-CoV-2 saltation events have a distinct mutational profile

We identify many residues along the genome that are repeatedly mutated in saltational branches (Table S1). To explore whether saltational branches display a distinct mutational signature, we evaluate whether a residue is more likely to have a nonsynonymous mutation in saltational branches compared with non-saltational branches. Across the genome, we find many residues that are more likely to be mutated on saltational branches than on non-saltational ones (Figure 3A). These sites concentrate in Spike (Figure 3B), particularly in the receptor-binding domain (RBD) (Figure 3C). These include many functionally relevant residues, known to affect immune escape (S:484 is mutated in 51 saltational branches, OR: 6.2 [95% CI: 4.6–8.2]), receptor binding (S:501 is mutated in 23 saltational branches, OR: 15.7 [95% CI: 9.8–24.2]) or viral entry (S:681 is mutated in 33 saltational branches, OR: 7.5 [95% CI: 5.1–10.6]). We also detect a high concentration of positions that are more likely to be mutated in saltational branches in the Nsp3 region of the ORF1a polyprotein (Figure 3D). This distinct mutational profile includes a higher frequency of mutations at known antigenic sites, including positions associated with antibody escape (Figure 3E) and surface accessible sites (Figure 3F), with nonsynonymous mutations in Spike enriched at surface residues (OR: 1.57 [95%CI: 1.43-1.73], Table S2). This suggests that shared selective pressures, such as immune escape, may act across saltational events and leave a detectable signature.

**Figure 3:**
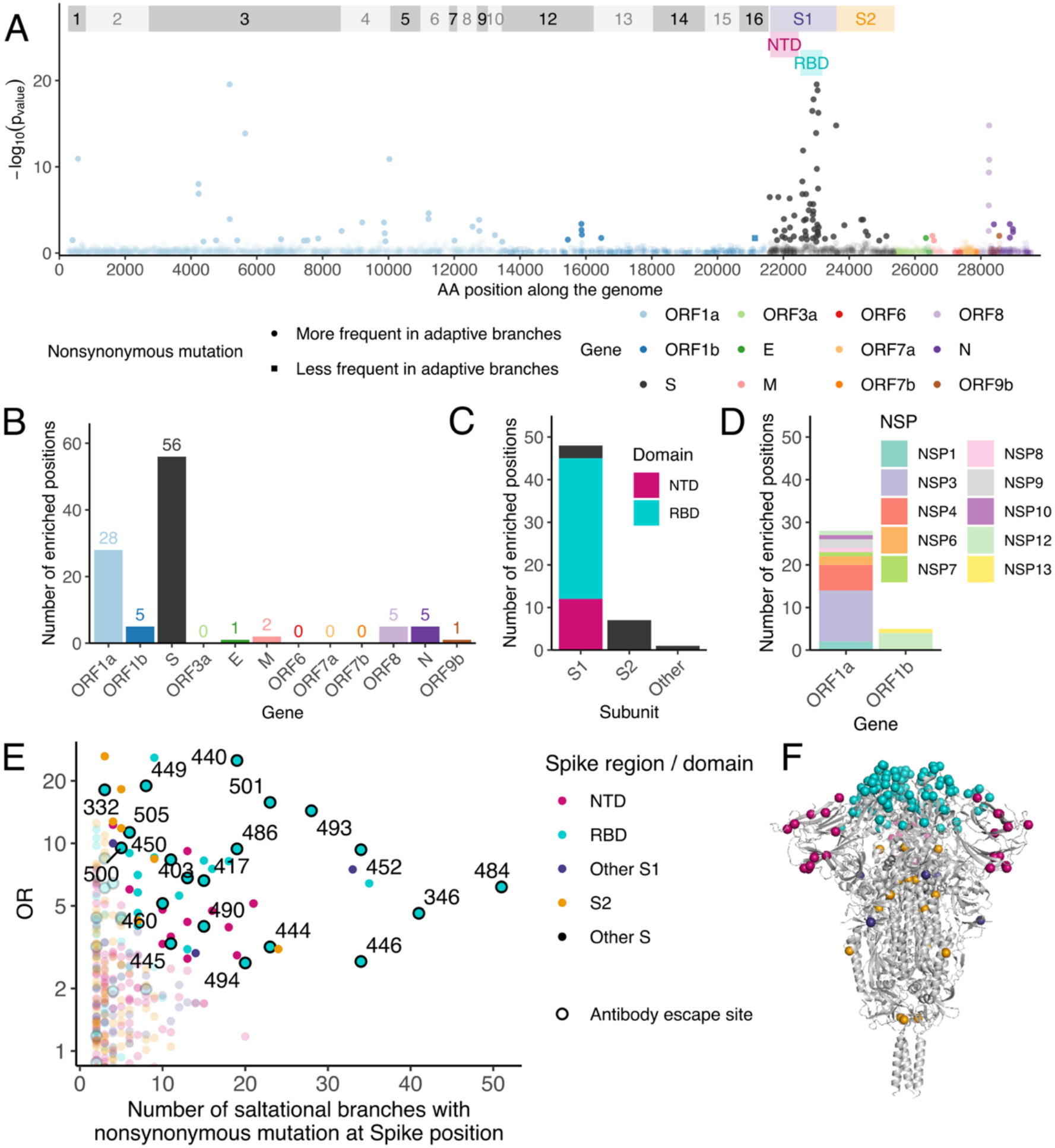
Mutational signature of SARS-CoV-2 saltational events. **A**. Manhattan plot showing p-values testing whether amino-acid residues are more or less frequently mutated on saltational branches than on non-saltational ones. P-values are adjusted for multiple testing (Benjamini-Hochberg). Non-significant positions (adjusted p-value greater than 0.05) are shown with greater transparency. Alternating grey shadings indicate the positions of the different Nsps and colored rectangles indicated Spike subunits and domains. Number of positions more likely to be mutated in saltational branches by **B**. gene, **C**. within Spike and **D**. within ORF1ab. **E**. Odds ratio for a nonsynonymous mutation occurring at a given residue within Spike on saltational branches compared with non-saltational ones, as a function of the number of saltational branches carrying a mutation at that residue. Residues that don’t reach statistical significance (adjusted p-value greater than 0.05) are shown with greater transparency. Points are circled if the residue has been associated with antibody escape^18–22^. **F**. Structure of the SARS-CoV-2 Spike protein (PDB: 7KRQ^23^), with residues colored if they are more likely to carry nonsynonymous mutations on saltational branches.

### The mutational signature of SARS-CoV-2 saltational events overlaps with that observed during persistent infections

The mutation enrichment profile of saltational branches may thus provide insights into the mechanisms that give rise to saltational events. To investigate this, we identify amino-acid mutations that occur significantly more often on saltational branches than on non-saltational ones (Table S3) and compare them with mutations reported to recur during persistent human infections, as well as after spillover into mink and deer populations since persistent infections and adaptation within other species were two hypotheses potentially explaining VOC-defining saltational branches (Figure 4A). This allows us to assess whether proposed mechanisms for the emergence of saltational events^7,11,12^ have recurrent mutations that are characteristic of SARS-CoV-2 saltational evolution. We find a strong overlap between mutations enriched in saltational branches and mutations that occur repeatedly during persistent infections: 26 enriched mutations (found at high frequency in saltational branches) overlap with mutations recurring in persistent infections, corresponding to 36% of the recurrent mutations identified in persistent infections. This includes mutations outside the Spike protein such as ORF1a:1638I (in 26 saltational branches), ORF1a:1795Q (in 14 saltational branches) or E:30I (in 7 saltational branches), that have been reported in persistent infections but that are observed at very low frequency in global sequences^13–16^. By contrast, only one enriched mutation overlaps with recurrent deer-associated mutations (2% of the deer recurrent mutation list we use), and 2 overlap with mink-associated mutations (33% of the mink recurrent mutation list). Moreover, mutations found to be recurring in mink occurred only nine times on saltational branches (Figure S7). Therefore, even though 2 out of 6 of the mink associated mutations are enriched on saltational branches, their absolute contribution to the mutation burden of saltational branches remains very limited. This overlap suggests a role for evolution within chronically infected individuals in the emergence of saltational events, though adaptation to other host species may also contribute to a small proportion of saltational events.

**Figure 4:**
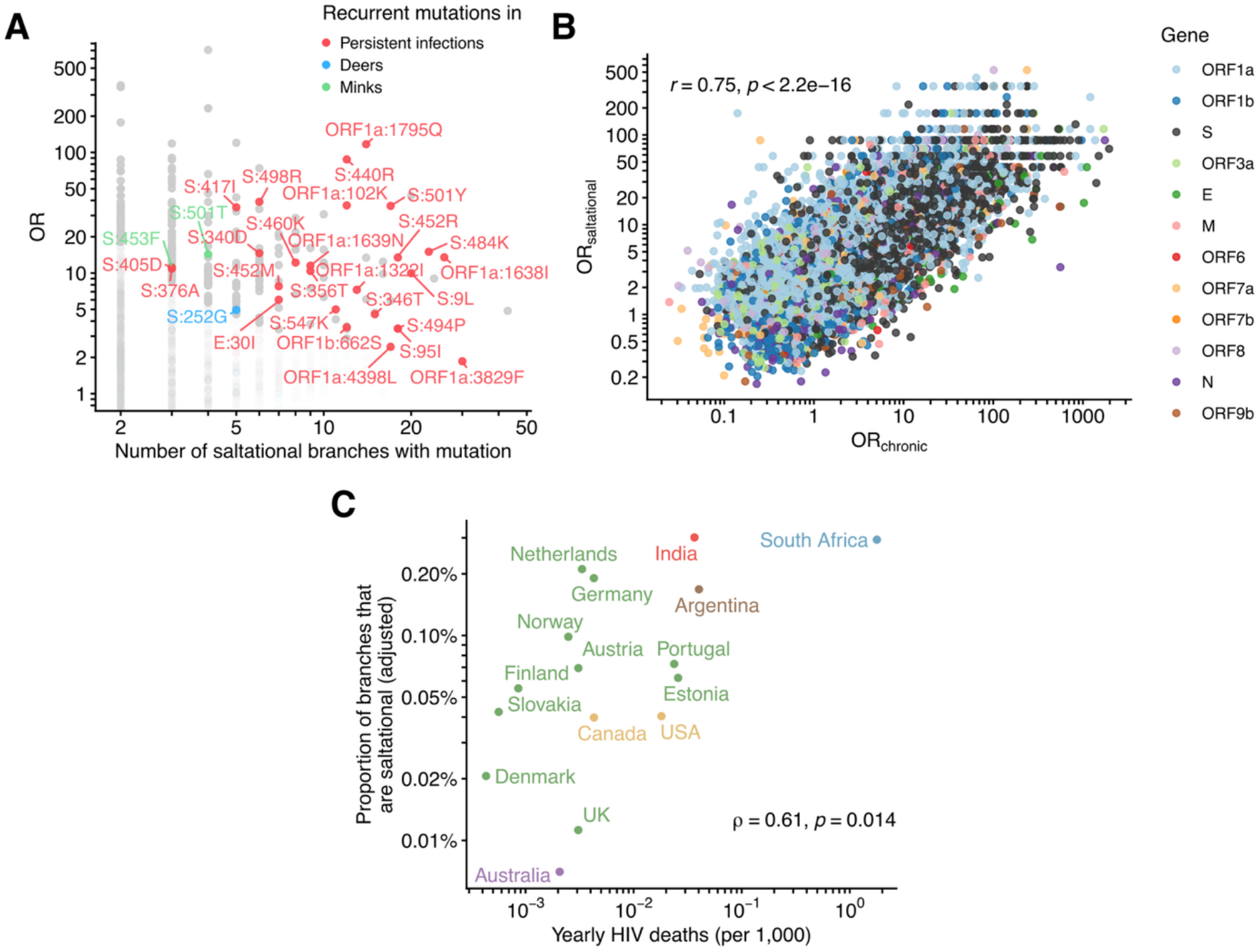
The mutational signature of SARS-CoV-2 saltational evolution supports the role of persistent human infections in their emergence. **A**. Odds ratio for a mutation occurring on saltational branches compared with non-saltational ones, as a function of the number of saltational branches carrying that mutation. Mutations that don’t reach statistical significance (adjusted p-value greater than 0.05) are shown with greater transparency. Points are colored according to whether the mutation has been reported to recur in persistent infections, deer or minks. **B**. Relationship between the odds ratio of a mutation occurring in saltational branches compared with non-saltational ones as a function of the odds ratio of the same mutation being observed in sequences from chronically infected individuals compared with a high-quality background sequence dataset^17^. **C**. Proportion of branches that are saltational as a function of HIV deaths (per 1,000 inhabitants) estimated for 2022^24,25^. To better visualize the trend in C, variables are displayed on a logarithmic scale and proportions are adjusted using pseudo-counts (Laplace smoothing) to enable displaying the proportion of branches that are saltational in Australia (equal to 0) (see Methods). The Spearman correlation coefficient is computed based on the unadjusted proportion.

Next, we evaluate whether the mutational signature of saltational branches overlaps with that of chronic infections by comparing the odds ratio of mutations occurring on saltational vs non-saltational branches to the odds ratio of the same mutation occurring in sequences collected from chronic infections relative to other circulating SARS-CoV-2 sequences^17^ (Figure 4B). We find a strong positive correlation of log odds ratio (Pearson *r = 0*.*75, p < 2*.*2⋅10*^*-16*^) between these two metrics, further supporting persistent infections as a likely source of many saltational events. This strong positive association remains observed when restricting the analysis to internal or terminal saltational branches (Figure S8) and when varying the thresholds used to define saltational branches (Figure S9).

Persistent SARS-CoV-2 infections have been repeatedly reported in immunocompromised individuals^7^, which suggests that the prevalence of conditions associated with immunosuppression may impact the likelihood of saltational events. Among these conditions, advanced human immunodeficiency virus (HIV) infection is of particular interest as it has been associated with prolonged time to SARS-CoV-2 clearance^6,26^, and heterogeneity in immune suppression levels that may be associated with some residual immunity against SARS-CoV-2 that could provide enough selection pressure to generate immune escape mutations^7,27^. Moreover, several VOCs (including Beta and Omicron) were first detected in regions experiencing a high HIV burden, which has led to the hypothesis that HIV may be linked to the emergence of highly mutated variants^7^. Here, we examine the relationship between spatial variation in the frequency of saltational events and heterogeneity in HIV burden across countries (Figure 4C). We find a moderate positive correlation between the country-level proportion of branches that are saltational and yearly AIDS mortality estimates (Spearman *ρ* = 0.60, *p* = 0.015). This association is reduced but remains consistent when excluding South Africa from the analysis (Spearman *ρ* = 0.52, *p* = 0.045). This association may also reflect regional differences in the prevalence of other immunosuppressive conditions, sampling strategies, infection burden or vaccination coverage. After adjusting for gross domestic product (GDP) per capita, the correlation remains positive but is no longer significant (Partial Spearman *ρ* =0.50, *p* = 0.060). Furthermore, this ecological analysis is based on only 16 countries which limits our ability to conclusively identify strong evidence for a relationship between the frequency of saltational branches and untreated HIV burden. This analysis is thus consistent with a relationship between untreated HIV burden and saltational evolution frequency but doesn’t provide conclusive evidence.

Overall, these results support the role of persistent human infections in the emergence of saltational events.

### Evidence of onward spread of a subset of saltational branches

So far, we have characterized the frequency and likely origin of saltational events. We next investigate their epidemiological consequences and the factors associated with their potential spread. Of the saltational events we identify, 70% [95% CI: 68-73%] occur on terminal branches (Figure 5A) and thus have no detected descendants in the phylogeny. Comparing the distribution of the number of detected descendants between non-saltational and saltational branches, we find that saltational branches are associated with a greater number of detected descendants, particularly in the upper tail of the distribution, but not with a greater probability of having any detected descendants (Figure S10). Whether a saltational event occurs on a terminal or internal branch however doesn’t directly translate to whether that event seeded a transmission cluster as descendants may not be sampled and (ii) some descendant cluster may reflect repeated sampling of the same individual^28^. To gain insights into saltational branches that seeded transmission clusters, we look for evidence of descendants with distinct characteristics (age, sex and geography). We find evidence consistent with spread to distinct individuals in 215 saltational branches (14%, 95% CI: 12-16%), with a subset spreading across borders: 67 spread to multiple countries (4.4%, 95% CI: 3.4-5.5%) and 53 to multiple continents (3.5%, 95% CI: 2.6-4.5%) (Figure 5B). Given incomplete sequencing and partial metadata availability, these proportions don’t directly measure the probability that saltational branches seed onward transmission and rather indicate that, for example, at least 14% of branches display evidence of transmission to distinct individuals. Among saltational branches with descendants, most remain detectable only briefly (median duration of 9 days), whereas a small fraction has descendants detectable over longer periods (90^th^ percentile: 103 days) (Figure S11). We find that the proportion of saltational branches with detected descendants increases when considering stricter thresholds for the definition of saltational branches (Figure S12). While the number of saltational branches identified varies considerably with the thresholds used (from 53 to 30,125), the proportion of saltational branches with detected descendants remains in a relatively similar range (from 29% to 43%).

**Figure 5:**
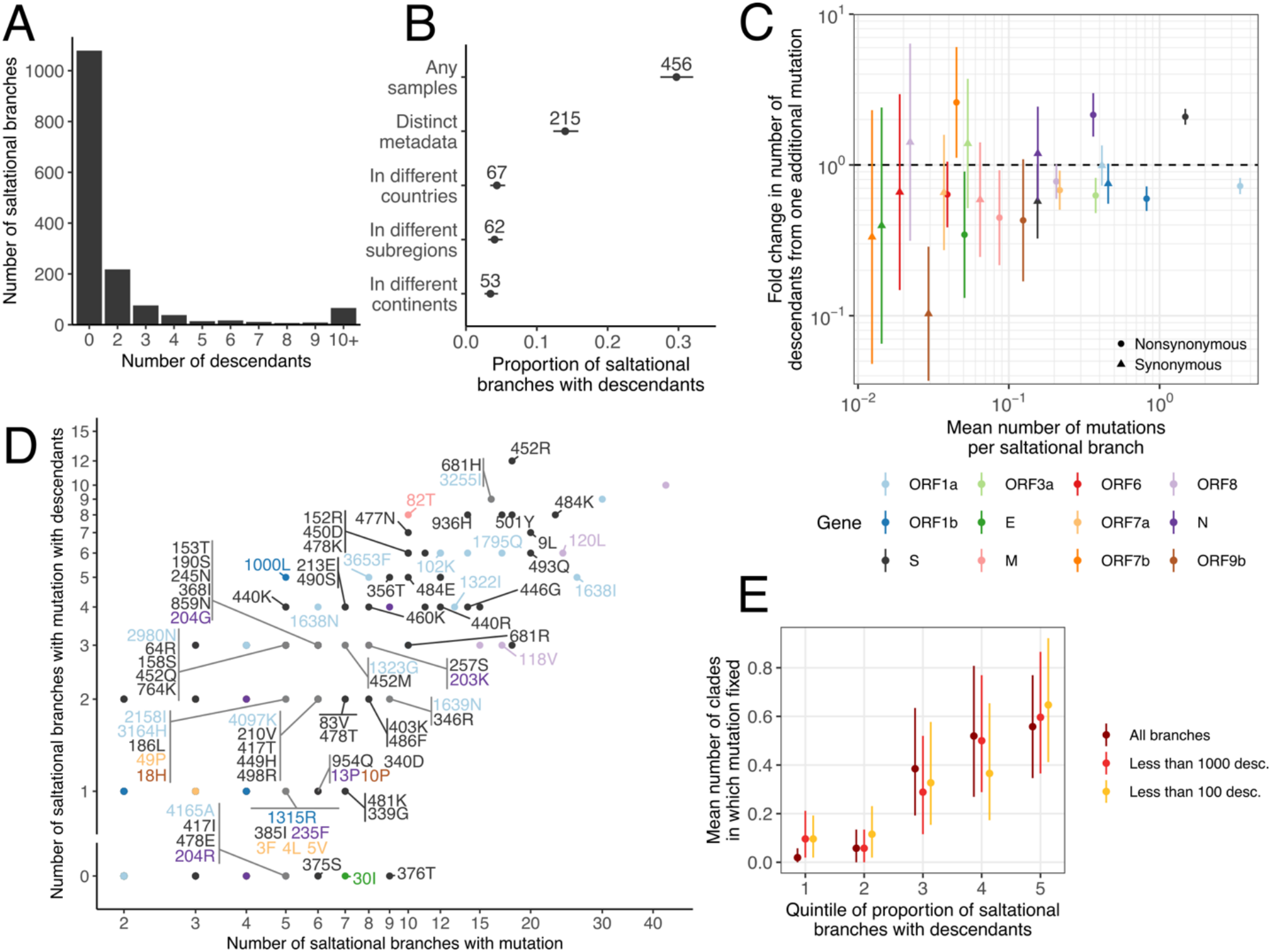
Patterns of onward spread from saltational branches. **A**. Distribution of descendant counts across saltational branches. **B**. Proportion of saltational branches with descendants across different spread criteria. Segments indicate 95% confidence intervals. **C**. Estimated fold-change in the number of descendants from one additional mutation as a function of the mean number of mutations per saltational branch and by mutation type (synonymous and nonsynonymous) and gene. Vertical segments indicate 95% confidence intervals. **D**. Number of saltational branches that have descendants and a mutation of interest as a function of the number of saltational branches that have that mutation. For clarity, we only write the labels of mutations with an OR of mutation occurring in saltational branches greater than 5 and that occur at least 5 times in saltational branches. **E**. Mean number of clades in which a mutation fixed at the global level by quintile of proportion of saltational branches with that mutation that have descendants. Vertical segments indicate 95% bootstrap confidence intervals. In E, we report the results based on all saltational branches or only saltational branches with less than 1000 or 100 descendants.

We explore whether the mutational composition of saltational branches is associated with the number of descendants each branch harbors. We model descendants counts as a function of synonymous and nonsynonymous mutation counts in each gene using a negative binomial regression (see Methods). Figure 5C depicts, for each gene and mutation type, the fold-change in expected descendant count associated with one additional mutation, as a function of the frequency of this mutation type in saltational branches. We find that one additional nonsynonymous mutation in spike is associated with a 2.1-fold [95% CI: 1.9-2.4] change in expected descendant counts, while one additional mutation in ORF1a and ORF1b are associated with a 0.72-fold [95% CI: 0.64-0.82] and a 0.60-fold [95% CI: 0.49-0.72] change in expected descendant counts. We identify other significant associations, including a positive effect of nonsynonymous mutations in N and ORF7b, but these mutations are relatively rare on saltational branches and are hence less likely to explain a large fraction of the overall variation in descendant counts.

Finally, we examine whether specific individual mutations are associated with saltational events leaving descendants (Figure 5D). We identify several mutations that occur repeatedly on saltational branches with descendants. By contrast, there are other mutations frequently observed on saltational branches that never leave detectable descendants. This suggests that the mutational composition of saltational branches may influence their ability to spread. Interestingly, S:484K, S:452R, S:681H and ORF1a:3255I, which recur frequently on saltational branches producing descendants, all are lineage-defining mutations in at least two emerging VOCs or Omicron sublineages. In comparison, we observe E:30I and the reversion S:376T on 7 and 9 saltational branches respectively, without any detectable descendants, and neither mutation was observed in successful SARS-CoV-2 clades. To quantify these associations, we compute the odds ratio for a saltational branch of having detected descendants according to whether it carries a mutation of interest (Table S4). The estimated odds ratio for S:484K, S:452R, S:681H and ORF1a:3255I are 1.3 [95%CI: 0.5-3.2], 4.8 [1.7-15.8], 3.0 [1.1-8.8] and 3.1 [1.0-9.8] respectively. None of these associations remained significant when adjusting for multiple testing. Given the small number of saltational branches that leave detectable descendants, we might not have enough power to robustly identify mutation-specific associations. Finally, we find that mutations more frequently observed on saltational branches with descendants also fix in a greater number of clades globally (Figure 5E). This relationship persists when excluding branches with more than 100 descendants, indicating that this result is not driven solely by the identification of clade-defining branches as saltational events.

Overall, we find evidence of onward transmission from a subset of saltational events and identify recurrent mutations within them that may be associated with their ability to spread.

## Discussion

Saltational evolution with the emergence of highly fit SARS-CoV-2 variants has been observed multiple times during SARS-CoV-2 evolution in the human population. Our systematic investigation of SARS-CoV-2 saltational evolution reveals that such events are a recurring feature of SARS-CoV-2 evolution, occurring at a low but detectable rate since the beginning of the pandemic to at least mid-2024 where our study data ends. Their distinct mutational signature is consistent with a role of chronic human infection in their emergence. While most of these saltational events don’t display evidence for onward transmission, a subset spread widely.

We find that the mutational signature of SARS-CoV-2 saltational events is informative about the potential mechanism of their emergence. Persistent replication of SARS-CoV-2 in chronically infected individuals with weakened immune system has been proposed to provide an evolutionary setting well-suited for the emergence of highly divergent SARS-CoV-2 variants^6,7^. The strong agreement between the mutational signature of saltational events and that of chronic human SARS-CoV-2 infections supports the role of persistent infections in SARS-CoV-2 saltational evolution (Figure 4B). We also identify a concentration of mutations at antigenically relevant residues in the Spike protein, particularly within the receptor-binding domain (Figure 3, Table S1, S3). This pattern is consistent with evolution during chronic infections of immunocompromised individuals, where selection pressure from partial immunity^29^ and treatment administration (including monoclonal antibodies^18–20,22^) may favor the acquisition of immune escape mutations. By contrast, the mutational signature of saltational events is inconsistent with that expected from reverse zoonosis in animal reservoirs such as minks or deer (Figure 4A).

Temporal and geographic variation in the frequency of saltational events (Figure 2) may thus reflect differences in emergence opportunity, which could stem from differences in infection burden, in population immunity, in the prevalence of persistent infections, changes in treatment regimens, or potential lineage-specific differences in mutational tolerance^30^. Variations in the detectability of saltational events may also explain some of these variations, for example with differences in sequencing intensity, in the patient populations captured by surveillance strategies or sequence quality.

While most saltational branches don’t display evidence for onward transmission, we identify 218 saltational events with evidence of transmission between individuals, a subset of which also dispersed widely, with for example 67 spreading across country borders (Figure 5). The distinct mutation patterns observed between branches that spread and those that don’t suggest that some mutations impact the ability for saltational branches to transmit between hosts. This is further supported by the observation that mutations more frequently found in spreading branches also tend to be successful at the global level. Interestingly, the highly divergent BA.3.2 lineage that was first detected after the end of our study period was characterized by a particularly high number of amino-acid changes in Spike, many at residues that we found to be more frequently mutated in saltational branches with descendants. These include S:9L and S:440R, two changes that we observed repeatedly on saltational branches with descendants but that were not seen in earlier variants of concern. This suggests that mutations recurring on saltational branches with detected descendants may help identify mutations that could later arise in globally successful variants, even if such mutations weren’t observed on previously successful variants. Overall, our findings are consistent with saltational events arising from persistent infections that select for viruses better adapted for persisting and replicating within hosts, but most of these variants are not well suited for transmission between hosts. However, a subset can transmit between hosts^31^, including some with potentially important epidemiological implications^3,4,32^. This may reflect pressure to escape immunity, either from the administration of treatment such as monoclonal antibodies, or from residual immunity, that may confer fitness advantages both at the within and between host level^7^.

Interestingly, although other pathogens can also cause persistent infections in immunocompromised individuals, a recurrent contribution of saltational events to their evolution hasn’t been demonstrated^33,34^. One possible explanation is that mutations selected within hosts may not confer an advantage or may even be incompatible with transmission between hosts^7^. However, for some pathogens including influenza, recurrent mutations arising during chronic infections in immunocompromised individuals overlap with mutations successful at the global level^34^, suggesting that similar processes may also occur in those systems. Yet influenza is densely surveilled, and its phylogenies accumulate mutations in a stepwise manner without the long, heavily mutated branches that define SARS-CoV-2 saltational evolution, suggesting this contrast reflects a genuine difference between the two viruses. That said, it remains possible that persistent infections contribute to global influenza evolution, even if the contribution appears on the surface weaker than for SARS-CoV-2.

Given the high potential risk of SARS-CoV-2 evolution during persistent infections, there are both individual and collective benefits associated with detecting and clearing such infections to mitigate the risk of emergence and transmission of new variants that may have major epidemiological consequences^6^. Given persistent infections have been disproportionately reported in immunocompromised patients^16^, efforts to reduce the burden of immunosuppressive conditions, when possible, may indirectly prevent the emergence of highly mutated SARS-CoV-2 variants by decreasing emergence opportunity. Advanced HIV has been associated with chronic SARS-CoV-2 infections during which highly divergent viruses can evolve^26,35,36^, and we find a moderate positive correlation between country-level HIV burden and the proportion of branches that are saltational (Figure 4C). In this context, recent disruptions to global HIV care^37,38^, including reduced access to antiretroviral therapy, could increase the number of individuals at high risk of persistent SARS-CoV-2 infections, thereby potentially amplifying opportunities for high-impact variant emergence. While we don’t aim to assess the impact of such disruptions on SARS-CoV-2 evolution, these considerations underline the interconnectedness of public health systems and crises, where the management of one epidemic may have indirect, here evolutionary, consequences for another, well beyond the initially affected populations.

Our ability to identify saltational events depends on the dataset analyzed and the criteria used to define them. Tightening or loosening these criteria would lead to detecting more or less events throughout the pandemic (Figure S2). Here, we thus don’t estimate the total number of saltational events that occurred throughout the study period. Instead, we report the subset of events that we can detect under a conservative criterion from a large high-quality sequence dataset. Our estimate of 0.03% of branches that are saltational is thus conservative and would increase when considering a less stringent definition for saltational evolution (Figure S2). Here, we classify individual branches in a large phylogeny as consistent with saltational evolution. Because the Viridian tree is built using UShER and does not encode indels^39^, our analysis of saltational evolution focuses on substitution patterns and doesn’t account for insertions and deletions. In practice, some saltational events may be distributed across several consecutive branches, for example because a single infection can span several branches^40^ or because of artifactual recombinant placements^41^ around long branches, which could decrease our power to detect them by diluting the signal. Finally, this study is enabled by the considerable scale of SARS-CoV-2 genomic surveillance globally^42^, along with the development of software tools able to handle large sequence datasets^39,43^ and efforts to improve sequence quality by reprocessing sequencing reads to correct for systematic errors^8,44,45^. Notably, our first attempts to perform this analysis on uncorrected publicly available consensus sequences^39^ mainly led to the identification of outlier branches containing mutations consistent with systematic errors occurring during genome assembly^8^.

To conclude, this work shows that saltational events are a recurrent component of SARS-CoV-2 evolution, linked to within-host evolutionary processes. Because a subset of these events seed onward human-to-human transmission, detecting and treating persistent SARS-CoV-2 infections is a tractable lever to mitigate the emergence of highly mutated, high-burden variants.

## Data and code availability

Data and code are available at https://github.com/blab/ncov-saltational. We directly deposited the files generated when analyzing the sequence data on Figshare to facilitate reproducing figures^46^.

## Acknowledgments

We thank Jesse Bloom and Sheri Harari for helpful discussions, and Victor Lin for support in downloading metadata from the ENA and SRA. We gratefully acknowledge the researchers and data contributors who collected the specimens, generated and deposited the raw sequence data and metadata into the European Nucleotide Archive (ENA) and Sequence Read Archive (SRA). We also thank the researchers behind the Viridian project for their work in improving the quality of publicly available SARS-CoV-2 sequence data and making this valuable resource available to the community. Analysis of chronic infections used data from GISAID^47,48^. We gratefully acknowledge all data contributors, i.e., the Authors and their Originating laboratories responsible for obtaining the specimens, and their Submitting laboratories for generating the genetic sequence and metadata and sharing via the GISAID Initiative, on which this research is based. The findings about the mutational signature of chronic infections of this study are based on 10,845,376 sequences accessible via https://doi.org/10.55876/gis8.260618hy. The supplemental table is available at https://github.com/blab/ncov-saltational.

## Author contributions

CTK, KK and TB conceived the study. CTK developed the methods. CTK, KK and RH analyzed the data. CTK wrote the first draft. All authors reviewed and edited the manuscript.

## Declaration of AI use

Generative AI tools were used to assist with software development and to support language editing of the manuscript.

## Methods

### Data

#### Large SARS-CoV-2 mutation-annotated tree of high-quality sequences with associated metadata

We rely on a publicly available high-quality mutation-annotated tree based on 4,471,579 consensus genomes that were reprocessed in an amplicon-aware manner using the Viridian tool^8^, which was developed to correct for systematic errors in genome assembly. We download corresponding associated run metadata from the SRA (geographic location, collection date and isolation source). We exclude samples with isolation sources that were consistent with environmental sampling, non-human hosts, non-respiratory clinical samples or laboratory experimental studies. We also only retain one sequencing run (and hence sequence) per biosample ID. We filter the Viridian tree using the *matUtils* library^49^. We then process the filtered mutation-annotated tree using the *BTE* package^50^ to extract child-parent relationships, the number of synonymous and nonsynonymous mutations occurring on each branch and whether these mutations are coded, the number of children and descendants for each node and the number of synonymous and nonsynonymous sites (defined as the number of possible synonymous and nonsynonymous mutations available per gene) at each node.

#### Recurring mutations in persistent infections, deer and mink infections

We use mutations that have been reported to recur in studies of persistent human SARS-CoV-2 infections (Table S5)^14–16,28,51^ as well as SARS-CoV-2 infections in minks (Table S6)^52–54^ and deer (Table S7)^53,55,56^.

We follow the approach described in Hisner et al.^17^ to identify sequences from individuals that are likely chronically infected and characterize the mutational signature of SARS-CoV-2 chronic human infections. Briefly, candidate chronic infections are manually identified from metadata associated with sequences deposited on GISAID^47,48^, unusually long branches in the phylogeny or the presence of mutations highly indicative of chronic infections. This approach led to the identification of 3843 sequences from distinct individuals that are likely chronically infected between April 2022 and January 2025. We compare the mutations that occur in these likely persistent infections to other circulating sequences to characterize the mutation profile of chronic infections (see below).

#### Country-level HIV mortality estimates and demographic covariates

To approximate advanced HIV disease burden, we use AIDS mortality estimates from the Global Burden of Disease for year 2022^24^. We convert these estimates to HIV deaths per 1,000 inhabitants using population size estimates from the World Bank for that year^25^. We also use per capita Gross Domestic Product estimates from the World Bank for that year^57^.

### Estimating branch-level *d*_*N*_*/d*_*S*_ from a mutation annotated tree

#### Approach

We aim to identify branches under positive selection by estimating the ratio of the number of nonsynonymous mutations per site to the number of synonymous mutations per site (*d*_*N*_*/d*_*S*_) for each branch and gene in a mutation-annotated tree. A dedicated approach is required for two reasons. First, most branches are short, and therefore contain few mutations, so reliable inference requires explicit uncertainty quantification and accounting for sparse mutation counts. Second, the size of the dataset requires a method that remains computationally tractable on a mutation-annotated tree containing millions of branches.

#### Notation

The index *i* refers to branches in the phylogeny and superscript *g* to the gene on which we infer the *d*_*N*_*/d*_*S*_ ratio. To estimate this quantity, we rely on the number of nonsynonymous and synonymous mutations (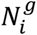 and 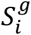) on branch *i* within gene *g*, and the number of nonsynonymous and synonymous sites (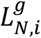 and 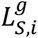) at the beginning of branch *i* within gene *g*. For each branch and gene, we introduce three parameters: (i) the number of nonsynonymous mutations per site 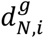, (ii) the number of synonymous mutations per site 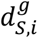 and (iii) the ratio between these quantities 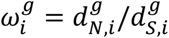.

#### Likelihood of the data

We assume that 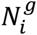 and 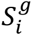 follow Poisson distributions parametrized as:

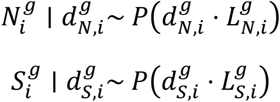

#### Prior distributions

We assume that the prior of the number of synonymous and nonsynonymous mutation per site are Gamma distributed:

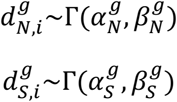

We can derive the prior distribution of 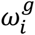 as the ratio of two independent Gamma distributions:

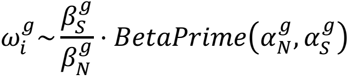

We set the prior means for the number of nonsynonymous and synonymous mutation per site in segment *g* to the corresponding tree-wide empirical ratios:

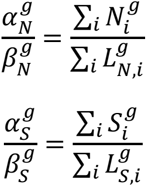

We explore a range of values for the coefficient of variation (CV) of the number of nonsynonymous and synonymous mutations per site and use posterior predictive checks (see below) to identify prior parametrizations that best reproduce the observed data.

#### Posterior distribution of parameters

The Gamma distribution is a conjugate prior of the Poisson distribution with exposure. We thus derive the posterior distribution of model parameters as:

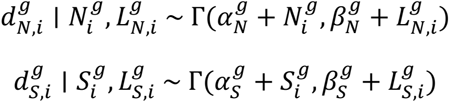

We deduce the posterior distribution of 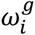 as the ratio of two independent Gamma distributions:

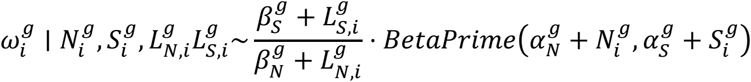

#### Posterior predictive checks

We perform posterior predictive checks (Figure S13) by drawing new observed counts of nonsynonymous and synonymous mutations 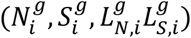 from:

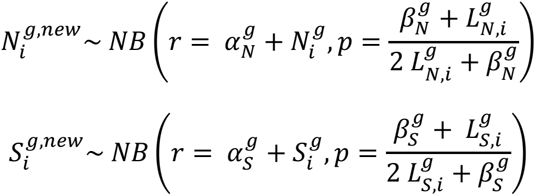

We compute the median number of branches 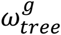 with each combination of nonsynonymous and synonymous mutations (*N, S*) across posterior simulations within gene *g*. We determine the coefficient of variation for the prior that maximizes the Pearson correlation between 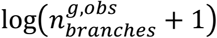. and 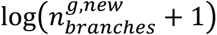. pooled across genes and (*N, S*) combinations, where 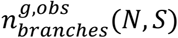 is the observed number of branches with *N* nonsynonymous and *S* synonymous mutations in gene *g* across the phylogeny (Figure S14-S15) (CV of 1.8).

#### Identifying branches characterized by d_N_/d_S_ > 1 at the gene level

To identify branches characterized by *d*_*N*_*/d*_*S*_ > 1 on gene *g*, we compute the Bayes Factor 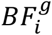 between the two competing hypotheses 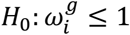 and 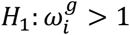 as the ratio of posterior and prior odds for 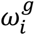 being greater than 1:

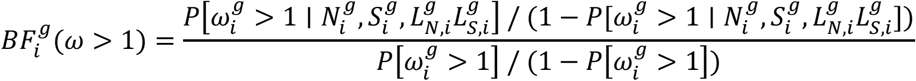

The posterior and prior probabilities 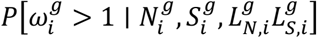 and 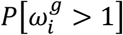 can be computed as:

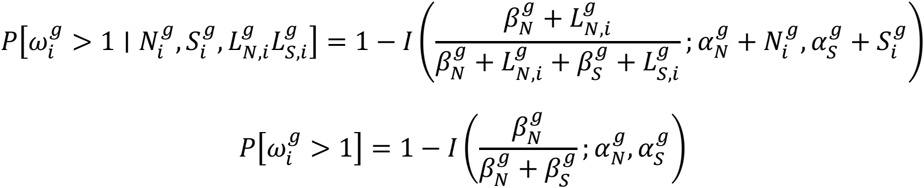

where *I*(*x*; *a, b*) is the regularized incomplete beta function evaluated in *x* with coefficients *a* and *b*. We classify branches as characterized by 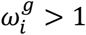 if 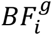 is above 20, interpreted as strong evidence for model *H*_1_ under the Kass-Raftery scale^9^.

#### Identifying branches characterized by genome-wide d_N_/d_S_ > 1

To assess evidence for positive selection at the genome level, we define branch-specific genome-wide number of nonsynonymous and synonymous mutations per site (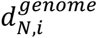 and 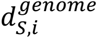), and the corresponding genome-wide ratio as:

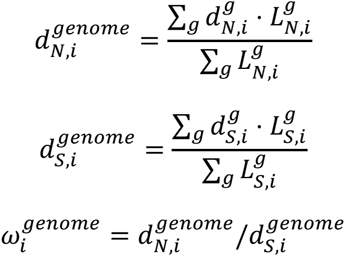

We use a Monte Carlo approach to explore the posterior distribution of 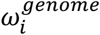. To do so, we draw *M* = 10,000 samples from each of the gene-specific posterior distributions of 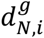 and 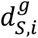. Using the formula above, we compute the corresponding *M* draws 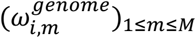 from the posterior distribution of 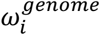. This enables us to approximate the posterior probability of 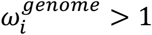 as:

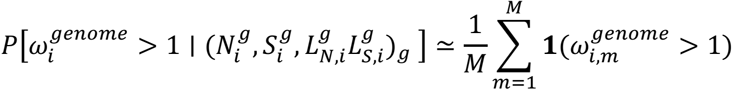

Similarly, we approximate the prior probability of 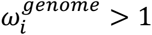 by drawing from the gene-specific prior distributions of the number of nonsynonymous and synonymous mutations per site. This enables us to compute 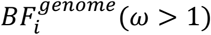 the ratio between the posterior and the prior odds of 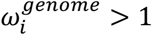.

To validate this genome-wide approach, we perform posterior predictive checks at the genome level by drawing synonymous and nonsynonymous mutation counts from the gene and branch-specific posterior predictive distributions and summing these counts across genes within each branch. Using the CV of 1.8 that maximizes the correlation between true and predicted number of branches at the gene level (see above) also provides a good predictive accuracy at the genome level (Pearson correlation coefficient of 0.9).

#### Compute time

Given the single gene model is conjugate, inference is fast and only required for the distinct combinations of synonymous and nonsynonymous mutation and site counts. For example, for the Spike model, there are 818 distinct combinations of synonymous and nonsynonymous mutation and site counts (distinct 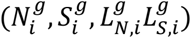) across the 5.3 million branches of the tree. Running the model for a single gene on the full tree thus took between 2 and 10 seconds using a single core of a personal machine (Apple M2 laptop). The genome-wide model requires Monte Carlo sampling for 777,909 mutational patterns (distinct combination of synonymous and nonsynonymous mutation and site counts across genes). It took 1 hour and 57 minutes using 6 cores on the same machine.

### Characterizing the signature of SARS-CoV-2 saltational evolution

#### Definition of saltational events

We define saltational events as single branches in the mutation-annotated tree with:

1. An unusually high number of nonsynonymous mutations (at least 5 at the genome level, corresponding to greater than the 99.5^th^ percentile)
2. Strong evidence for positive selection (measured by a Bayes Factor at the genome level or within a gene greater than 20 ^9^)

Stricter thresholds rapidly reduce the number of branches classified as saltational (Figure S2). Branches identified as saltational under less stringent criteria still differ from background branches, though with a weaker separation. We choose this conservative threshold combination (Bayes Factor above 20 and at least 5 nonsynonymous mutations) as a compromise between identifying a sufficiently large set of branches for detailed downstream analyses (∼1000) while ensuring a stringent definition of saltational evolution. We also report sensitivity analyses exploring the impact of varying the thresholds used to define saltational branches on the relationship between the mutational signature of saltational evolution and persistent human infections (Figure S9) and on the likelihood of saltational branches having detected descendants (Figure S12). To mitigate the impact of recombination, we exclude from potential saltational events branches annotated as leading to new recombining Pango lineages in the Viridian tree^8^.

#### Comparison with expectation under a null model

We compare the characteristics of the saltational branches we identify in the *Viridian* tree with the ones obtained under a null expectation. We simulate 20 null datasets by fixing the gene and branch-specific total mutation counts and redrawing mutation types (synonymous and nonsynonymous) using the tree-wide d_N_/d_S_ average for each gene 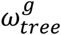. More specifically, we draw the number of nonsynonymous and synonymous mutations on branch *i* within gene *g* from a Binomial distribution:

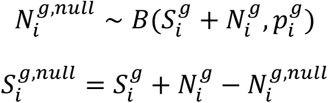

With

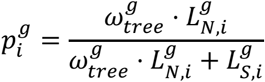

and

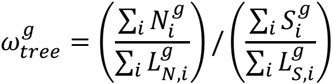

We then apply our saltational evolution criterion to detect saltational branches on these simulated datasets. We compare the characteristics of the observed saltational branches and saltational branches under the null, both in terms of number of branches and mutation characteristics (Figure S3). This null model doesn’t explore whether long branches with many mutations occur more frequently than one would expect given we keep mutation counts constant. However, it enables to explore whether some branches carry more nonsynonymous mutations than we would expect.

#### Temporal signature of saltational events

We define the year of each branch as the year of the node at the end of the branch furthest from the root. For internal nodes or terminal nodes with missing sample collection date information, we run Chronumental^43^ to infer node times. We compute the proportion of branches that are saltational each year and compute 95% Wilson confidence intervals around these proportions.

#### Geographical signature of saltational events

We determine internal nodes’ geographical locations based on the geographies of their descendants. If all descendants from a node come from the same country, we attribute the internal node to this country. Otherwise, we annotate this node as having “Multiple descendants”. We use the same approach to reconstruct subregions (defined using the United Nations geoscheme) and continents of internal nodes. We chose this coarse and conservative approach to geographical attribution for internal nodes as we were concerned substantial heterogeneities in sampling and sequencing intensity could bias ancestral state reconstruction. It also allows us to analyze the entire tree, without relying on subsampling.

#### Mutational signature of saltational events

To evaluate whether saltational branches have a distinct mutational profile, we use Fisher’s exact tests (two-sided) to assess whether nonsynonymous mutations are more likely to occur at specific amino acid positions in saltational branches compared with non-saltational branches. Using mutations as the statistical unit enables us to account for the higher number of mutations occurring on saltational branches (Figure 1D). By contrast, a branch-level analysis of whether a position is more likely to be mutated in saltational branches would be biased by the excess of mutations in saltational branches, which could lead to identifying positions as more mutated on saltational branches only because mutations occur more frequently on this type of branch. We identify 2,199 amino-acid positions that are mutated (nonsynonymously) at least twice in saltational branches. For each of these amino-acid positions, we run Fisher tests (two-sided) and report p-values adjusted for multiple testing (Benjamini-Hochberg correction). We use a Type I error threshold *α* of 0.05. We also perform Fisher’s exact tests (two-sided) to assess whether nonsynonymous mutations are more likely to occur on Spike surface accessible residues^58^ in saltational branches (Table S2).

To complement this position-level analysis, we assess whether specific amino-acid changes are more likely to occur on saltational branches (e.g. S:484K rather than any mutation occurring at position S:484). As not all mutations are possible on all branches (e.g. S:484K is not possible if a K is already present at position 484), we perform Fisher’s exact tests (two-sided) using only branches on which that mutation is possible (e.g. branches without a K at position 484 in that specific example). For a given mutation *m*, the odds ratio (OR) of occurrence on saltational branches compared with non-saltational ones is equal to:

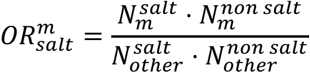

Here, 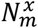 is the number of occurrences of mutation *m* on branches of type *x* (saltational or non-saltational) where the mutation is possible, and 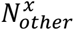 is the number of other mutations on branches of type *x* on which mutation *m* is possible.

#### Comparison of the mutational signature of saltational events with that of persistent infections

We next compare the log odds ratio of occurrence of mutations on saltational branches (vs non-saltational branches) with the log odds ratio of occurrence of mutations in chronic infections compared with background sequences^17^. As odds ratio can be null (if the mutation never occurs), we rely on a modified version of odds ratios:

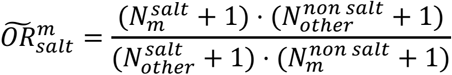

To compute the odds ratio of a mutation occurring in sequences from chronically infected individuals vs background sequences, we rely on a large manually assembled dataset containing 3843 sequences from distinct individuals that are likely chronically infected^17^. We compare the mutations that occur in the sequences from likely chronic infections to other circulating sequences as:

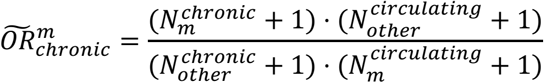

where 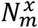 is the number of occurrences of mutation *m* on sequences of type *x* (chronic or other circulating, and 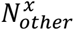 is the number of other mutations on sequences of type *x*. This dataset doesn’t report whether a mutation is possible (if the derived amino-acid differs from the ancestral one). However, we don’t expect this to impact the OR computation meaningfully as the OR of observing a mutation in saltational branches is little impacted by accounting for whether a nonsynonymous mutation is possible on a given branch (Figure S16).

To perform the comparison between 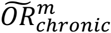 and 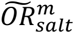, we focus on mutations that are observed at least once in either saltational branches or sequences from chronic infection (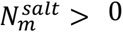 or 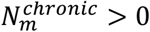), and that were observed at least once in both of the OR computations meaning:

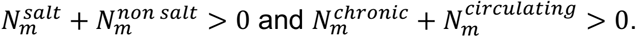

#### Quantifying saltational branches’ onward transmission patterns

We define descendants from saltational branches as terminal tips (corresponding to distinct collected samples) descending from the saltational branch. We determine a saltational branch to have evidence of spread to multiple distinct individuals if we can identify descendant samples with distinct metadata (either age, sex or geographic location information). Many samples don’t have fine-grained metadata information (age and sex) and it is hence not always possible to determine whether it spread to multiple distinct individuals. We also determine whether there is evidence for a saltational event to have spread across geographies (countries, subregions or continents) if descendant samples are collected in multiple geographies. Finally, we compare the distribution of the number of descendants between saltational branches and non-saltational branches.

We investigate whether the mutational composition of saltational branches is associated with the number of descendants from this saltational event. To do so, we model descendant counts as a function of the number of synonymous and nonsynonymous mutations in each gene. We define descendant counts as 0 for branches without detected descendants and as the number of terminal tips descending from that branch minus 1 otherwise. That way, our metric of descendant counts reflects the number of additional descendants observed beyond the initial saltational branch. We estimate the relationship between mutation counts and descendant counts using a negative binomial regression. More specifically, let *D*_*i*_ denote the number of descendants from saltational branch *i*. We model *D*_*i*_ as:

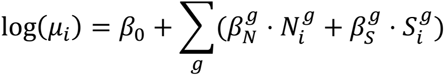

Where 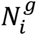 and 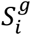 are respectively the number of nonsynonymous and synonymous mutations in gene *g* on branch *i* and 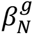 and 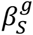 are parameters to be estimated. We fit the negative binomial regression in R using the *glmmTMB* package^59,60^. We decided to perform a negative binomial regression over a Poisson regression after performing a likelihood ratio test of the two models that yielded a p-value < 2.2⋅10^-16^ for the negative binomial model over the Poisson one, suggesting overdispersion in the data that needs to be accounted for. As a small number of saltational branches have a very high number of descendants, their inclusion in the analysis can impact parameter estimates (Figure S17). To mitigate the impact of a few branches with many descendants, we present estimates by restricting the analysis to saltational events that have less than 50,000 descendants in the global phylogeny.

#### Assessing the dynamics of mutations enriched in saltational branches at the between-host level

We compare the frequency at which mutations associated with saltational branches give rise to descendants with the frequency at which these mutations fix at the global level. We measure how often a mutation fixed throughout the pandemic by counting the number of non-recombinant Nextstrain clades in the open all-time Nextstrain public tree^61,62^ (downloaded on April 20, 2026) in which that mutation fixed. We only focus on mutations that are enriched in saltational branches. We consider that a mutation fixes within a clade if it is present in at least 80% of the tips descending from this clade and if the mutation arose on the branch leading to the clade or within the clade. We then group mutations into quintiles based on the proportion of saltational branches carrying that mutation that have descendants. For each quintile, we compute the mean number of clades in which these mutations fixed. We also compute uncertainty around the mean using 95% bootstrap confidence intervals from 2000 bootstrap draws. As some saltational branches with descendants are clade-defining, we perform a sensitivity analysis by removing saltational branches with more than 100 or 1000 descendants from the computations.

## References

1. Hill, V. et al. The origins and molecular evolution of SARS-CoV-2 lineage B.1.1.7 in the UK. Virus Evol. 8, veac080 (2022).

2. Tegally, H. et al. Detection of a SARS-CoV-2 variant of concern in South Africa. Nature 592, 438–443 (2021).

3. Faria, N. R. et al. Genomics and epidemiology of the P.1 SARS-CoV-2 lineage in Manaus, Brazil. Science 372, 815–821 (2021).

4. Viana, R. et al. Rapid epidemic expansion of the SARS-CoV-2 Omicron variant in southern Africa. Nature 603, 679–686 (2022).

5. Volz, E. Fitness, growth and transmissibility of SARS-CoV-2 genetic variants. Nat. Rev. Genet. 24, 724–734 (2023).

6. Machkovech, H. M. et al. Persistent SARS-CoV-2 infection: significance and implications. Lancet Infect. Dis. 24, e453–e462 (2024).

7. Sigal, A., Neher, R. A. & Lessells, R. J. The consequences of SARS-CoV-2 within-host persistence. Nat. Rev. Microbiol. 23, 288–302 (2025).

8. Hunt, M. et al. Addressing pandemic-wide systematic errors in the SARS-CoV-2 phylogeny. Nat. Methods 23, 653–662 (2026).

9. Kass, R. E. & Raftery, A. E. Bayes Factors. J. Am. Stat. Assoc. 90, 773 (1995).

10. Aksamentov, I., Roemer, C., Hodcroft, E. & Neher, R. Nextclade: clade assignment, mutation calling and quality control for viral genomes. J. Open Source Softw. 6, 3773 (2021).

11. Markov, P. V. et al. The evolution of SARS-CoV-2. Nat. Rev. Microbiol. 21, 361–379 (2023).

12. Carabelli, A. M. et al. SARS-CoV-2 variant biology: immune escape, transmission and fitness. Nat. Rev. Microbiol. 21, 162–177 (2023).

13. Velasquez-Reyes, J. M. et al. Characterisation of a persistent SARS-CoV-2 infection lasting more than 750 days in a person living with HIV: a genomic analysis. Lancet Microbe 6, 101122 (2025).

14. Ghafari, M. et al. Prevalence of persistent SARS-CoV-2 in a large community surveillance study. Nature 626, 1094–1101 (2024).

15. Wilkinson, S. A. J. et al. Recurrent SARS-CoV-2 mutations in immunodeficient patients. Virus Evol. 8, veac050 (2022).

16. Khurana, M. P. et al. Large-scale genomic surveillance reveals immunosuppression drives mutation dynamics in persistent SARS-CoV-2 infections. medRxiv (2025) doi:10.1101/2025.02.10.25321987.

17. Hisner, R. & Martin, D. P. Genetic evidence indicates the evolutionary importance of the SARS-CoV-2 ORF9b protein. bioRxiv (2026) doi:10.64898/2026.02.23.707522.

18. Starr, T. N. et al. Prospective mapping of viral mutations that escape antibodies used to treat COVID-19. Science 371, 850–854 (2021).

19. Starr, T. N., Greaney, A. J., Dingens, A. S. & Bloom, J. D. Complete map of SARS-CoV-2 RBD mutations that escape the monoclonal antibody LY-CoV555 and its cocktail with LY-CoV016. Cell Rep. Med. 2, 100255 (2021).

20. Dong, J. et al. Genetic and structural basis for SARS-CoV-2 variant neutralization by a two-antibody cocktail. Nat. Microbiol. 6, 1233–1244 (2021).

21. Greaney, A. J. et al. Mapping mutations to the SARS-CoV-2 RBD that escape binding by different classes of antibodies. Nat. Commun. 12, 4196 (2021).

22. Dadonaite, B. et al. Spike mutations that affect the function and antigenicity of recent KP.3.1.1-like SARS-CoV-2 variants. J. Virol. 99, e0142325 (2025).

23. Zhang, J. et al. Structural impact on SARS-CoV-2 spike protein by D614G substitution. Science 372, 525–530 (2021).

24. Global Burden of Disease Collaborative Network. Global Burden of Disease Study 2023 (GBD 2023). Seattle, United States: Institute for Health Metrics and Evaluation (IHME) (2025).

25. World Bank Open Data. World Bank Open Data https://data.worldbank.org/.

26. Karim, F. et al. Clearance of persistent SARS-CoV-2 associates with increased neutralizing antibodies in advanced HIV disease post-ART initiation. Nat. Commun. 15, 2360 (2024).

27. Raglow, Z. et al. SARS-CoV-2 shedding and evolution in patients who were immunocompromised during the omicron period: a multicentre, prospective analysis. Lancet Microbe 5, e235–e246 (2024).

28. Harari, S., Miller, D., Fleishon, S., Burstein, D. & Stern, A. Using big sequencing data to identify chronic SARS-Coronavirus-2 infections. Nat. Commun. 15, 648 (2024).

29. Grenfell, B. T. et al. Unifying the epidemiological and evolutionary dynamics of pathogens. Science 303, 327–332 (2004).

30. Sesta, L. & Neher, R. A. Epistasis and the changing fitness landscapes of SARS-CoV-2. bioRxiv (2026) doi:10.64898/2026.03.12.711354.

31. Gonzalez-Reiche, A. S. et al. Sequential intrahost evolution and onward transmission of SARS-CoV-2 variants. Nat. Commun. 14, 3235 (2023).

32. Volz, E. et al. Assessing transmissibility of SARS-CoV-2 lineage B.1.1.7 in England. Nature 593, 266–269 (2021).

33. Raglow, Z. & Lauring, A. S. Virus evolution in prolonged infections of immunocompromised individuals. Clin. Chem. 71, 109–118 (2025).

34. Xue, K. S. et al. Parallel evolution of influenza across multiple spatiotemporal scales. Elife 6, e26875 (2017).

35. García-Martínez de Artola, D. et al. A prolonged Delta SARS-CoV-2 infection during the Omicron wave in an HIV immunosuppressed patient. Int. J. Infect. Dis. 162, 108233 (2026).

36. Nabieva, E. et al. A highly divergent sample from a nearly extinct SARS-CoV-2 lineage in a patient with long-term COVID-19. Front. Cell. Infect. Microbiol. 15, 1623390 (2025).

37. Cavalcanti, D. M. et al. Evaluating the impact of two decades of USAID interventions and projecting the effects of defunding on mortality up to 2030: a retrospective impact evaluation and forecasting analysis. Lancet 406, 283–294 (2025).

38. Brink, D. T. et al. Impact of an international HIV funding crisis on HIV infections and mortality in low-income and middle-income countries: a modelling study. Lancet HIV 12, e346–e354 (2025).

39. Turakhia, Y. et al. Ultrafast Sample placement on Existing tRees (UShER) enables real-time phylogenetics for the SARS-CoV-2 pandemic. Nat. Genet. 53, 809–816 (2021).

40. Didelot, X., Fraser, C., Gardy, J. & Colijn, C. Genomic infectious disease epidemiology in partially sampled and ongoing outbreaks. Mol. Biol. Evol. msw075 (2017).

41. Turakhia, Y. et al. Pandemic-scale phylogenomics reveals the SARS-CoV-2 recombination landscape. Nature 609, 994–997 (2022).

42. Brito, A. F. et al. Global disparities in SARS-CoV-2 genomic surveillance. Nat. Commun. 13, 7003 (2022).

43. Sanderson, T. Chronumental: time tree estimation from very large phylogenies. bioRxiv (2021) doi:10.1101/2021.10.27.465994.

44. Turakhia, Y. et al. Stability of SARS-CoV-2 phylogenies. PLoS Genet. 16, e1009175 (2020).

45. De Maio, N. et al. Issues with SARS-CoV-2 sequencing data. virological.orghttps://virological.org/t/issues-with-sars-cov-2-sequencing-data/473 (2020).

46. Tran-Kiem, C. blab/ncov-saltational workflow outputs. figshare 10.6084/M9.FIGSHARE.32736657 (2026).

47. Shu, Y. & McCauley, J. GISAID: Global initiative on sharing all influenza data – from vision to reality. Euro Surveill. 22, (2017).

48. GISAID - gisaid.org. https://gisaid.org/.

49. McBroome, J. et al. A daily-updated database and tools for comprehensive SARS-CoV-2 mutation-annotated trees. Mol. Biol. Evol. 38, 5819–5824 (2021).

50. BTE — usher_wiki 0.0.2 documentation. https://usher-wiki.readthedocs.io/en/latest/bte.html.

51. Harari, S. et al. Drivers of adaptive evolution during chronic SARS-CoV-2 infections. Nat. Med. 28, 1501–1508 (2022).

52. Zhou, J. et al. Mutations that adapt SARS-CoV-2 to mink or ferret do not increase fitness in the human airway. Cell Rep. 38, 110344 (2022).

53. Tan, C. C. S. et al. Transmission of SARS-CoV-2 from humans to animals and potential host adaptation. Nat. Commun. 13, 2988 (2022).

54. Iglesias-Caballero, M. et al. Genomic context of SARS-CoV-2 outbreaks in farmed mink in Spain during pandemic: Unveiling host adaptation mechanisms. Int. J. Mol. Sci. 25, 5499 (2024).

55. Marques, A. D. et al. Evolution of SARS-CoV-2 in white-tailed deer in Pennsylvania 2021-2024. PLoS Pathog. 21, e1012883 (2025).

56. Feng, A. et al. Transmission of SARS-CoV-2 in free-ranging white-tailed deer in the United States. Nat. Commun. 14, 4078 (2023).

57. GDP per capita (current US$). World Bank Open Data https://data.worldbank.org/indicator/NY.GDP.PCAP.CD.

58. von Bülow, S., Sikora, M., Blanc, F. E. C., Covino, R. & Hummer, G. Antibody accessibility determines location of spike surface mutations in SARS-CoV-2 variants. PLoS Comput. Biol. 19, e1010822 (2023).

59. Brooks, M. et al. GlmmTMB balances speed and flexibility among packages for zero-inflated generalized linear mixed modeling. R J. 9, 378 (2017).

60. McGillycuddy, M., Warton, D. I., Popovic, G. & Bolker, B. M. Parsimoniously fitting large multivariate random effects in glmmTMB. J. Stat. Softw. 112, (2025).

61. nextstrain.org. Nextstrain SARS-CoV-2 all-time global public tree. https://nextstrain.org/ncov/open/global/all-time.

62. Hadfield, J. et al. Nextstrain: real-time tracking of pathogen evolution. Bioinformatics 34, 4121–4123 (2018).

