## Supplement for "SARS-CoV-2 saltational events are recurrent and trace to persistent human infections"

### Supplementary information for SARS-CoV-2 saltational events are recurrent and trace to persistent human infections


#### Supplementary figures

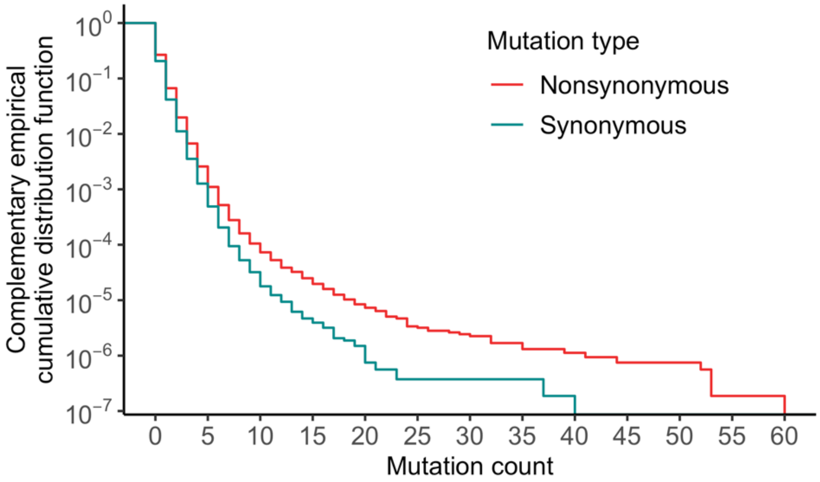


**Figure S1: Complementary empirical cumulative distribution function of mutation count across branches in the phylogeny and by mutation type (nonsynonymous or synonymous).** For each mutation count value on the x-axis, the lines indicate the proportion of branches that have at least that many mutations.

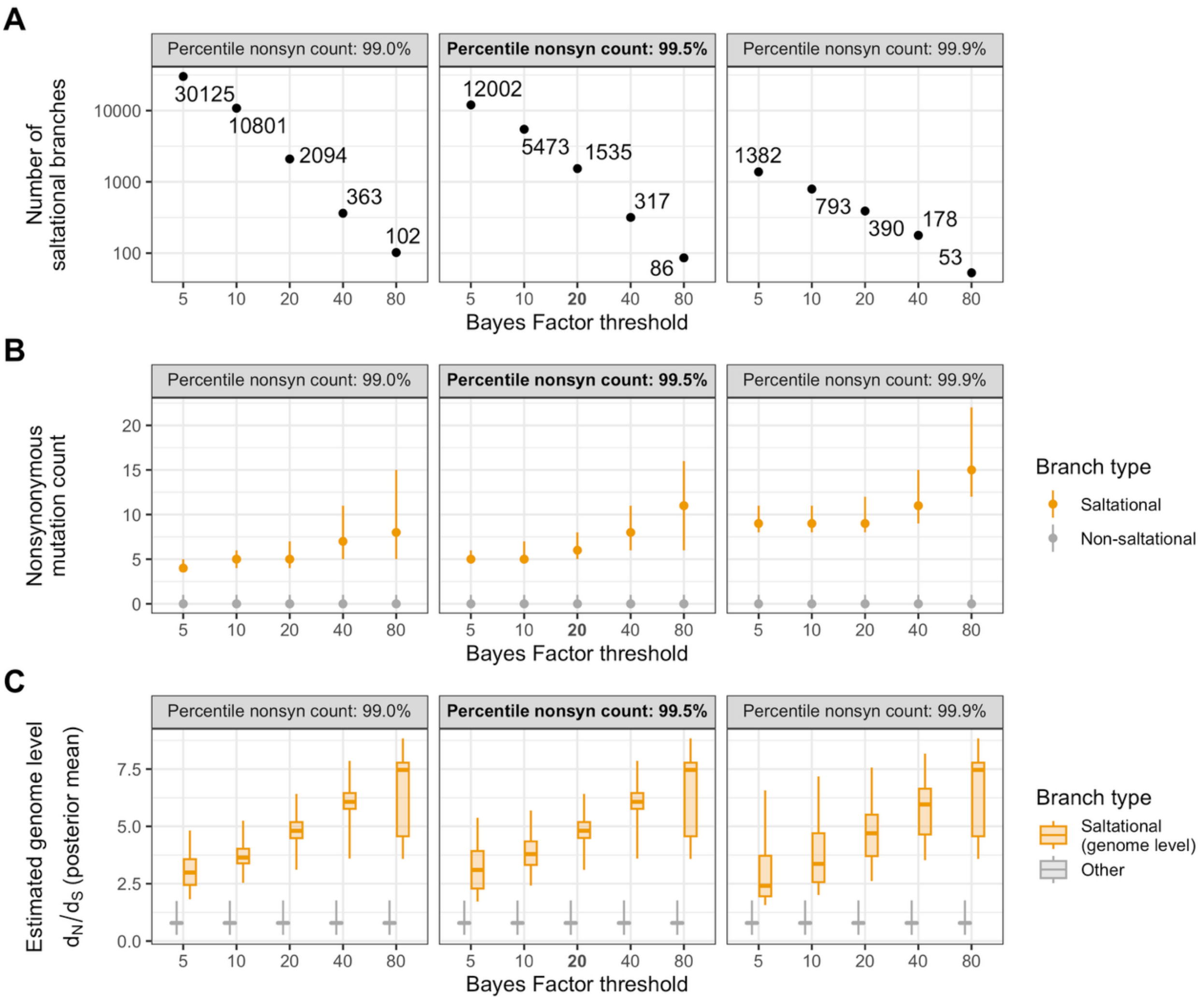


**Figure S2: Impact of saltational branch definition on the characteristics of saltational branches.** We explore the impact of varying the Bayes Factor threshold used to identify branches under positive selection (x-axis) and the minimum genome-wide nonsynonymous mutation count required for classification as saltational (facets). **A.** Number of branches classified as saltational under different criteria. **B.** Median nonsynonymous mutation count on saltational and non-saltational branches under different criteria. Vertical segments indicate interquartile ranges. **C.** Distribution of posterior mean genome-wide $d_N/d_S$ ratio for branches classified as saltational at the genome level and for other branches. Boxplots indicate 2.5th, 25th, 50th, 75th and 97.5th percentiles.

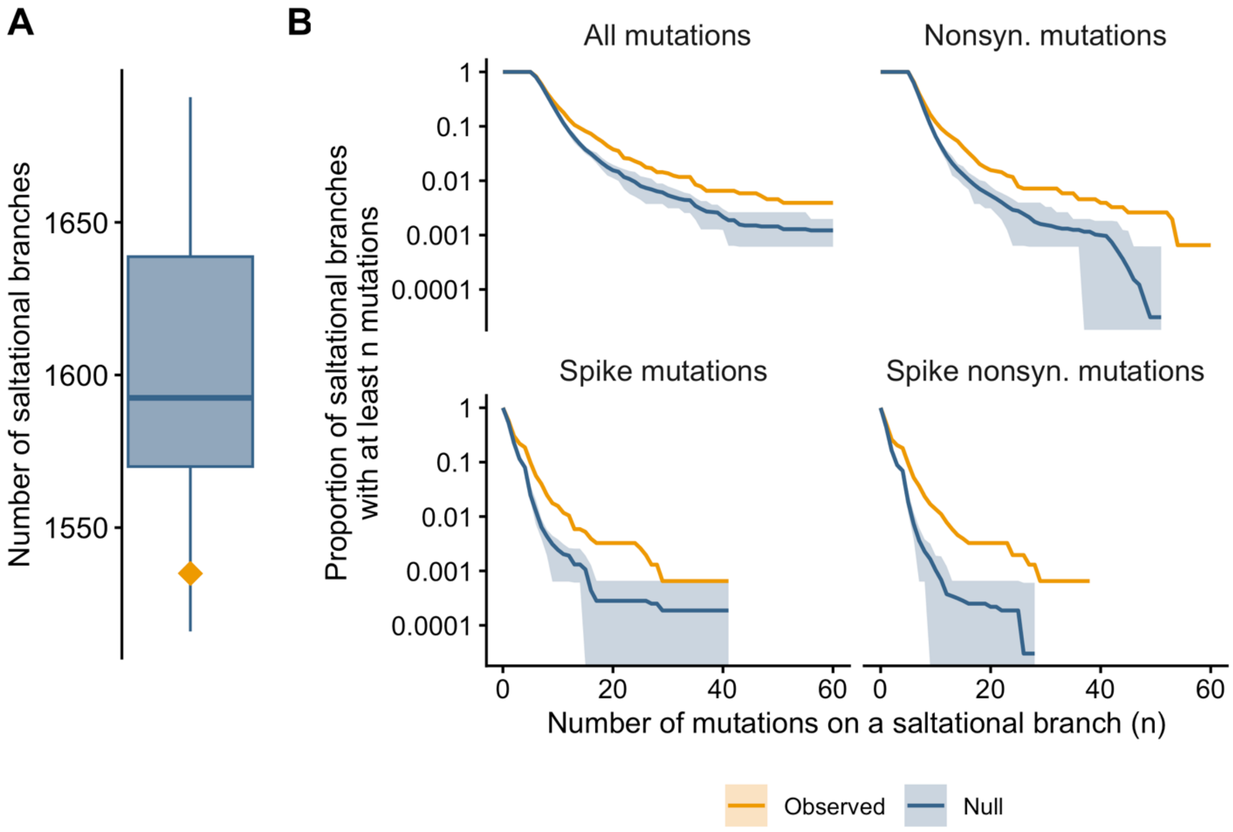


**Figure S3: Characteristics of observed saltational branches and branches identified as saltational under a null model preserving branch-specific substitution counts. A.** Number of branches classified as saltational from 20 simulated null datasets (boxplot). The orange diamond indicates the number of branches classified as saltational in the Viridian tree. **B.** Complementary cumulative distribution function of mutation counts on observed saltational branches (orange) and on saltational branches obtained from the null model (blue). In A, the boxplot indicates the interquartile range and the range across all simulations (whiskers). In B, the solid blue lines correspond to the mean across the 20 replicates run and the shaded ribbon corresponds to the range across simulations.

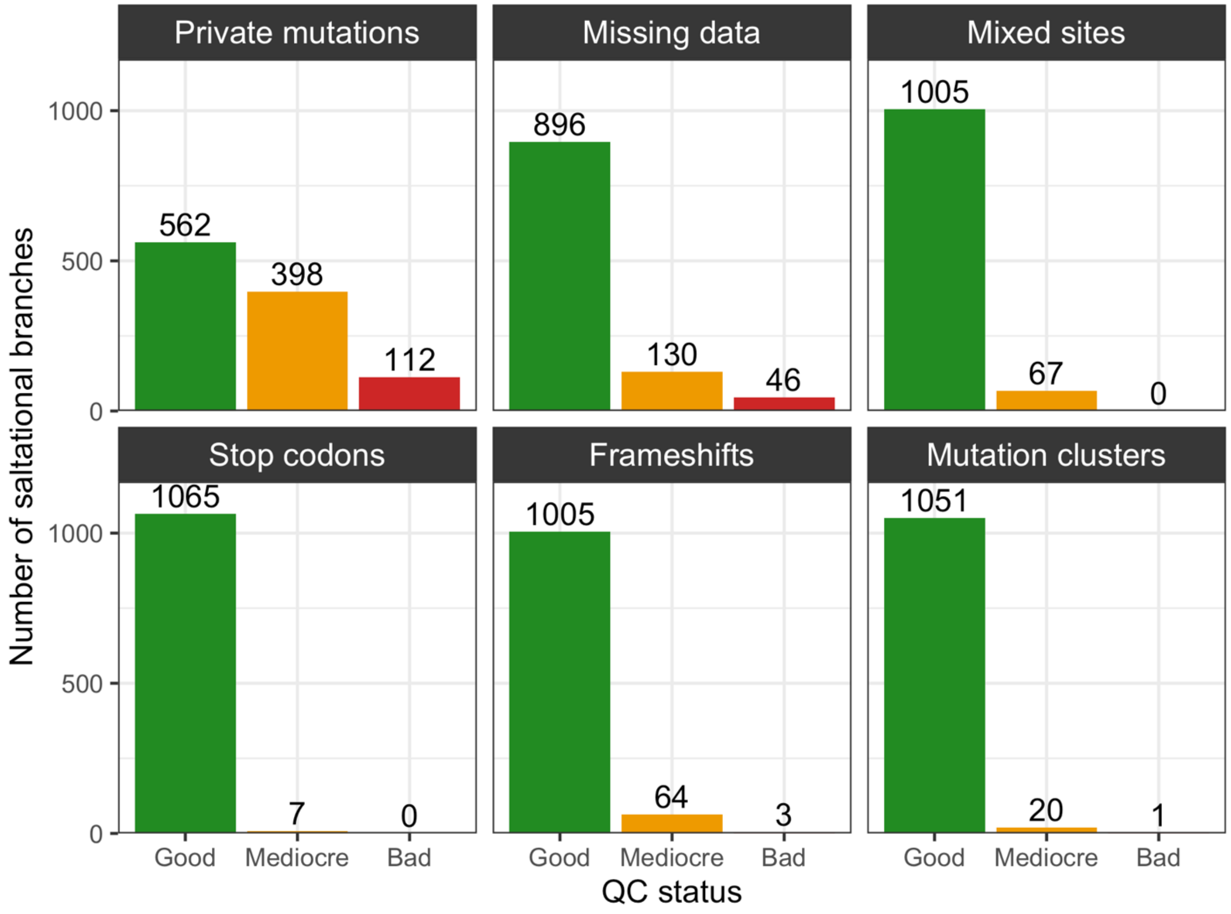


**Figure S4: Quality-control metrics for terminal saltational branches.** Distribution of Nextclade QC statuses (good, mediocre and bad) for each of the QC rules (private mutations missing data, mixed sites, stop codons, frameshifts and mutation clusters) among terminal saltational branches.

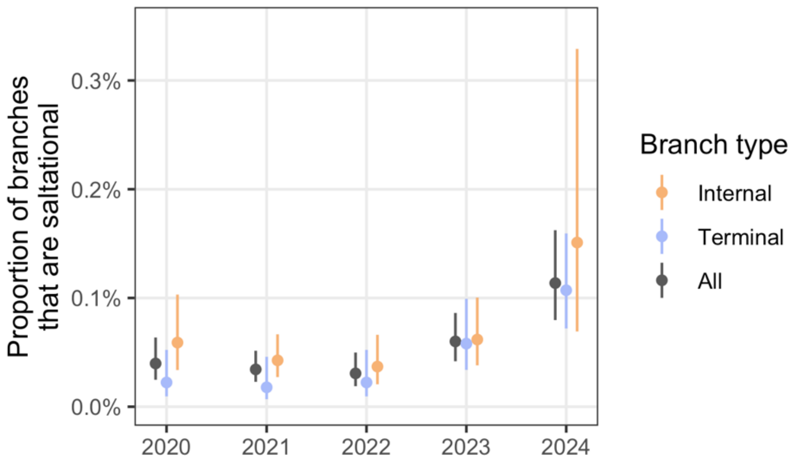


**Figure S5: Sensitivity analysis based on a subsampled tree with the same number of tips every year.** We downsampled the tree used in the analysis by randomly selecting, for each year, the same number of tips as available for 2025. The plot shows the proportion of branches classified as saltational by year.

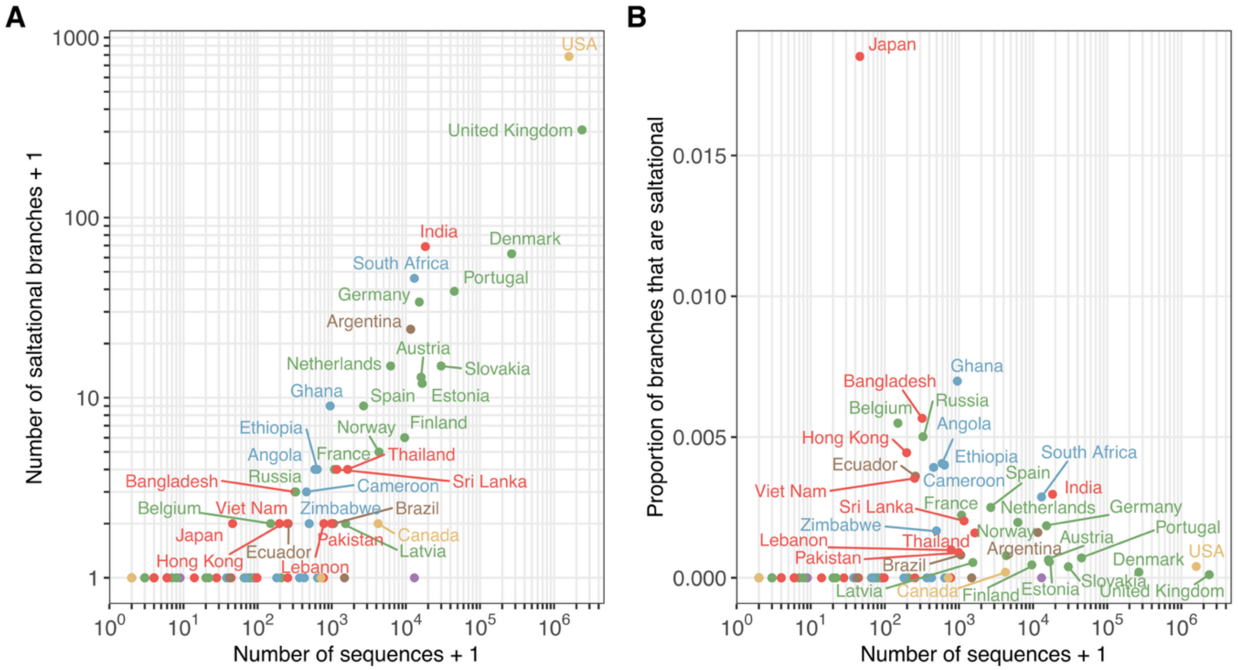


**Figure S6: Relationship between the number of saltational branches identified in a country and the number of sequences from this country included in this analysis. A.** Number of branches identified as saltational + 1 and **B.** proportion of branches that are saltational as a function of the number of sequences from that country.

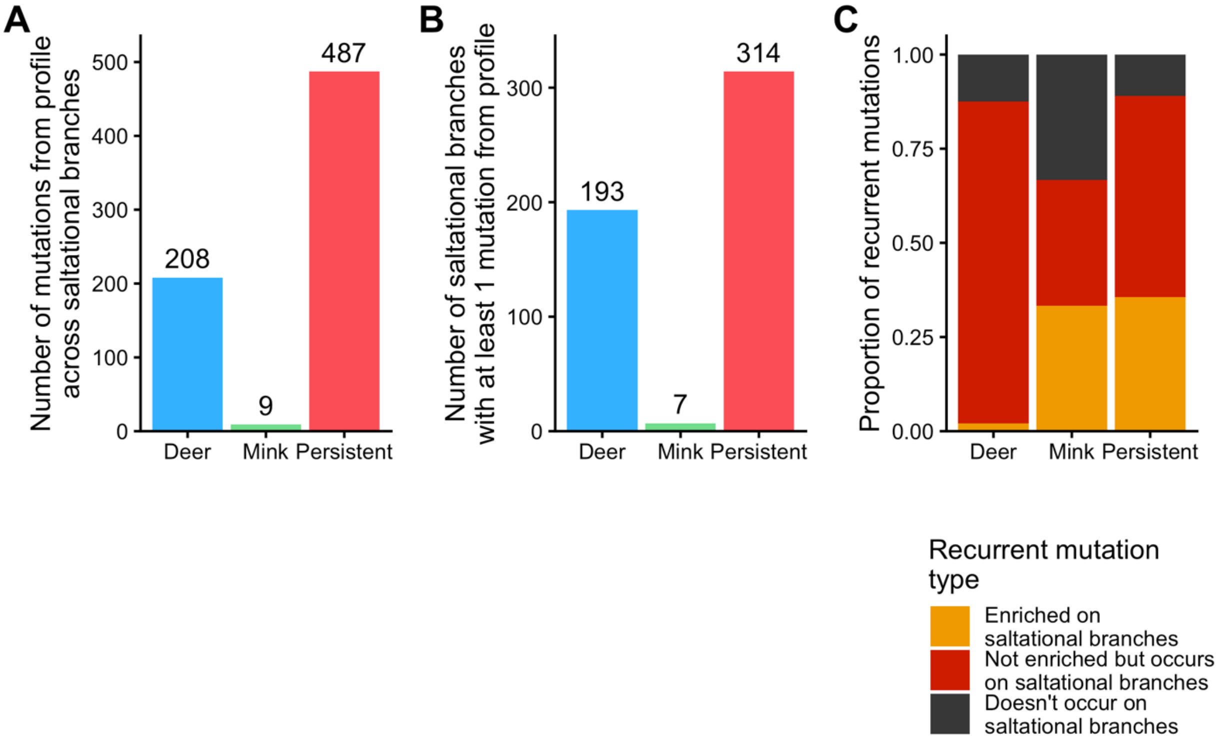


**Figure S7: Overlap between recurrent mutation lists and mutations observed on saltational branches. A.** Number of nonsynonymous mutations from the deer, mink and persistent infection lists (called profile) observed on saltational branches. **B.** Number of saltational branches carrying at least 1 mutation from the deer, mink and persistent infection recurrent mutation lists. **C.** Proportion of recurrent mutation from each profile that are enriched on saltational branches, observed but not enriched or not observed on saltational branches. Enrichment is based on p-values from Fisher tests adjusted from multiple testing (Benjamini-Hochberg correction) being lower than 0.05 (see Methods).

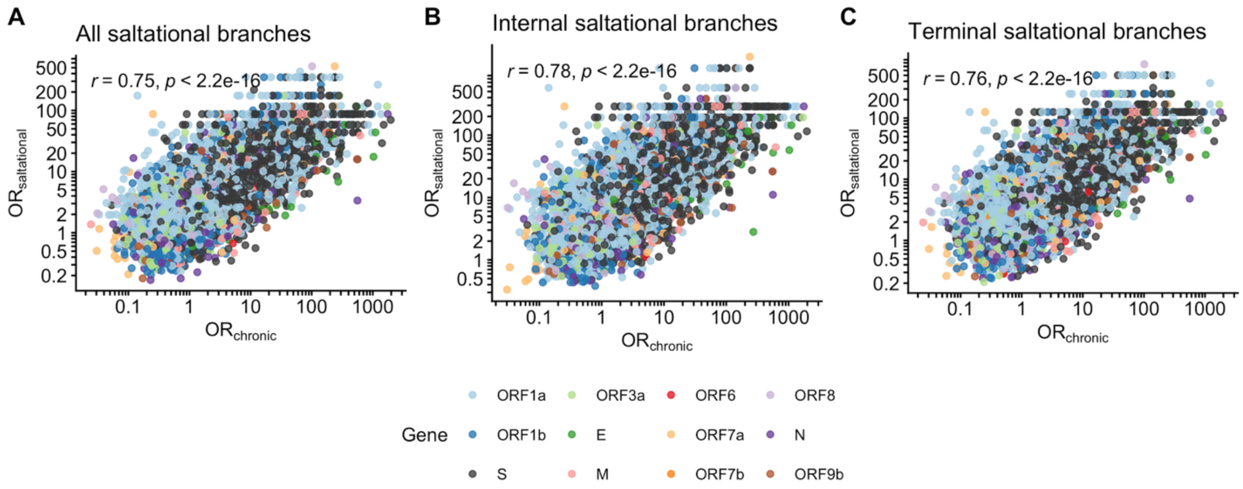


**Figure S8: Sensitivity analysis of the relationship between the mutational signature of saltational branches and that of human persistent infections.** Each panel shows the odds ratio of a mutation occurring in saltational branches compared with non-saltational branches as a function of the odds ratio of the same mutation being observed in sequences from chronically infected individuals compared with a high-quality background sequence dataset using **A.** all saltational branches, **B.** only internal saltational branches (with detected descendants) and **C.** only terminal saltational branches.

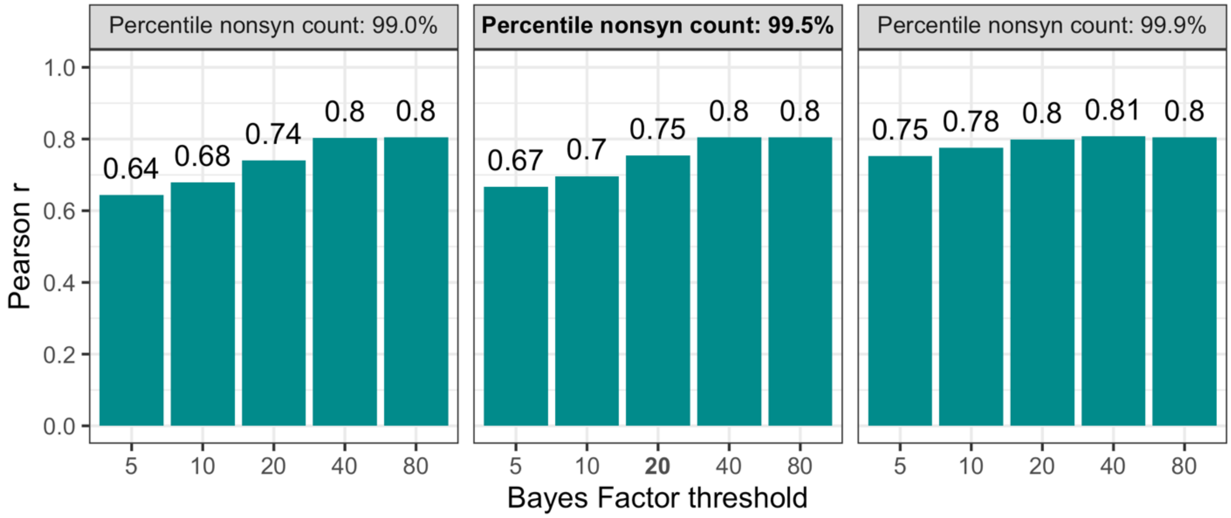


**Figure S9: Impact of saltational branch definition on the agreement between the mutational signature of saltational evolution and that of persistent human infections.** We explore the impact of varying the Bayes Factor threshold used to identify branches under positive selection (x-axis) and the minimum genome-wide nonsynonymous mutation count required for classification as saltational (facets) on the Pearson correlation coefficient between the log odds ratio of a nonsynonymous mutation occurring in saltational vs non-saltational branches and in persistent human infection vs high-quality background sequences.

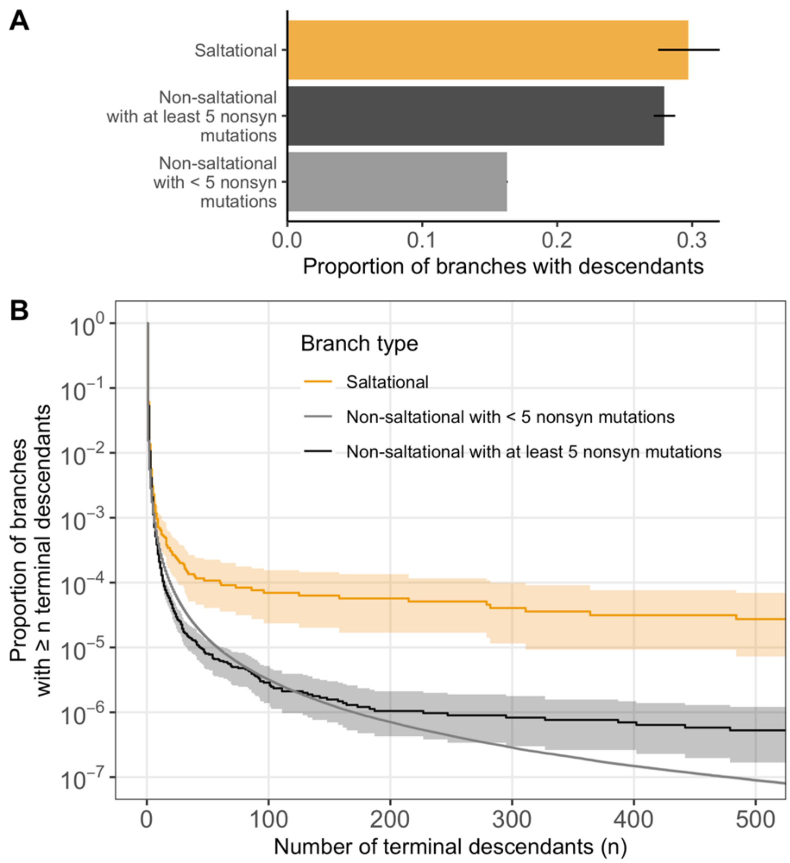


**Figure S10: Number of detected descendants per branch type. A.** Proportion of branches with detected descendants across saltational branches, non-saltational branches with at least 5 nonsynonymous mutations and non-saltational branches with less than 5 nonsynonymous mutations. Segments indicate 95% confidence intervals for proportions. **B.** Complementary cumulative distribution of the number of detected descendants for each branch type. The solid line indicates the proportion of branches with at least *n* detected descendants. Shaded areas indicate 95% bootstrap confidence intervals obtained by resampling branches with replacement within each branch type. We used 10,000 bootstrap replicates for saltational branches and non-saltational branches with at least 5 nonsynonymous mutations and 1,000 bootstrap replicates for non-saltational branches with less than 5 nonsynonymous mutations (as computational requirements were larger for the latter).

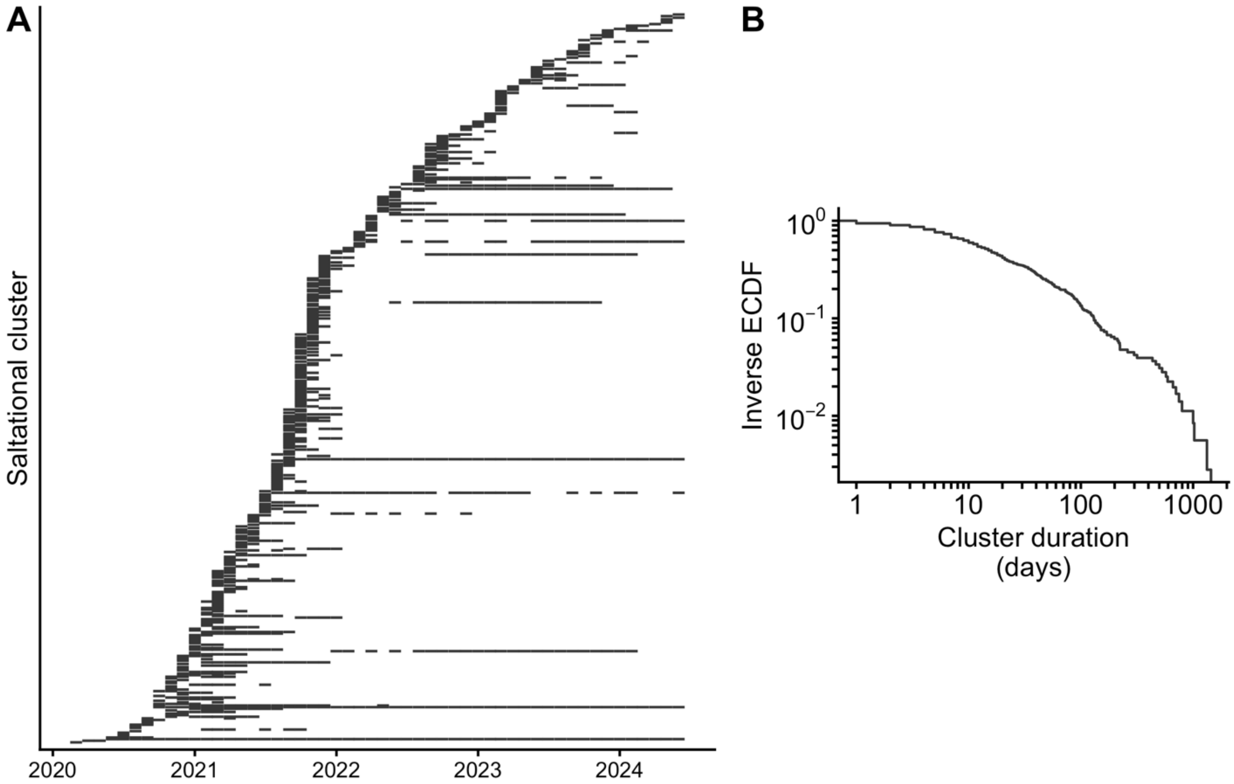


**Figure S11: Persistence of clusters descending from saltational events. A.** Temporal window during which clusters descending from saltational events are observed. Each line on the y-axis corresponds to a cluster descending from saltational events. Clusters are ordered by inferred saltational branch date. A colored tile indicates that this cluster has sequences collected during the corresponding month. **B.** Inverse empirical cumulative distribution function (ECDF) of cluster durations, measured as the delay between the sequence collection date of the latest and earliest sequence descending from this saltational branch.

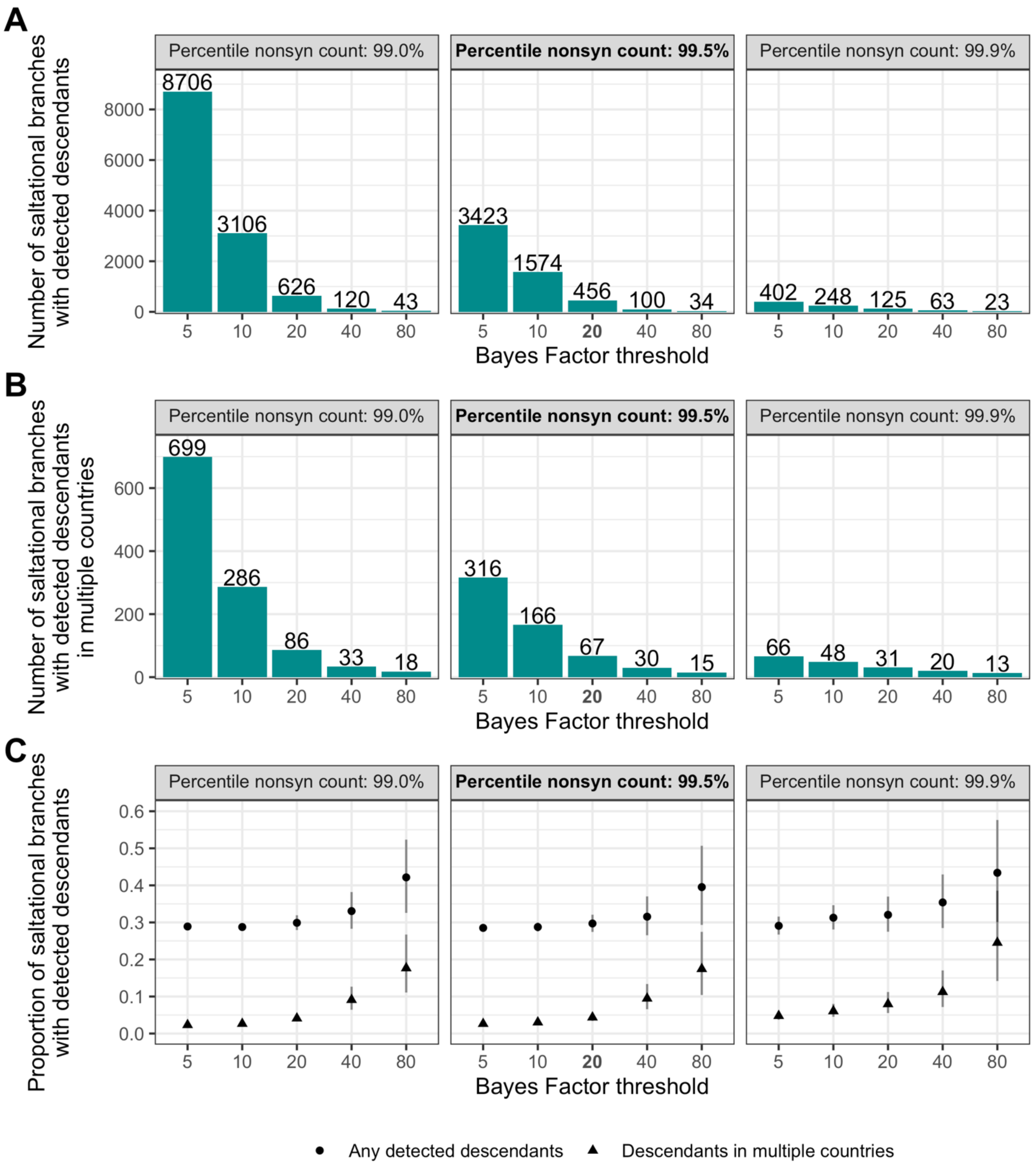


**Figure S12: Impact of saltational branch definition on detected descendants' patterns across saltational branches.** We explore the impact of varying the Bayes Factor threshold used to identify branches under positive selection (x-axis) and the minimum genome-wide nonsynonymous mutation count required for classification as saltational (facets). **A.** Number of saltational branches with detected descendants under different criteria. **B.** Number of saltational branches with detected descendants in multiple countries under different criteria. **C.** Proportion of saltational branches with detected descendants. Segments indicate 95% confidence intervals.

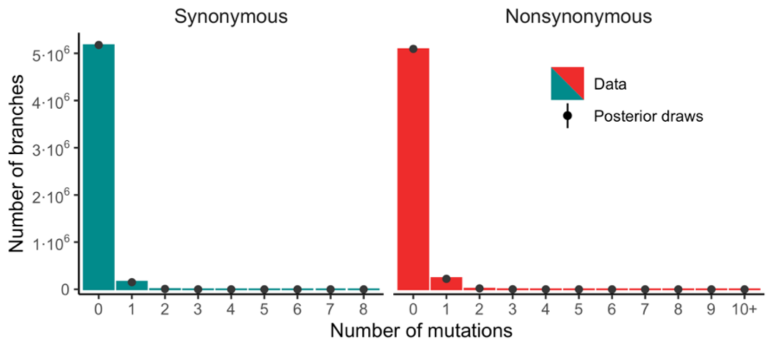


**Figure S13: Distribution of synonymous** (left) **and nonsynonymous** (right) **mutation counts on Spike by branch**. Bars depict observed counts in the large SARS-CoV-2 phylogeny. Points and vertical segments depict median and 95% credible intervals from posterior draws.

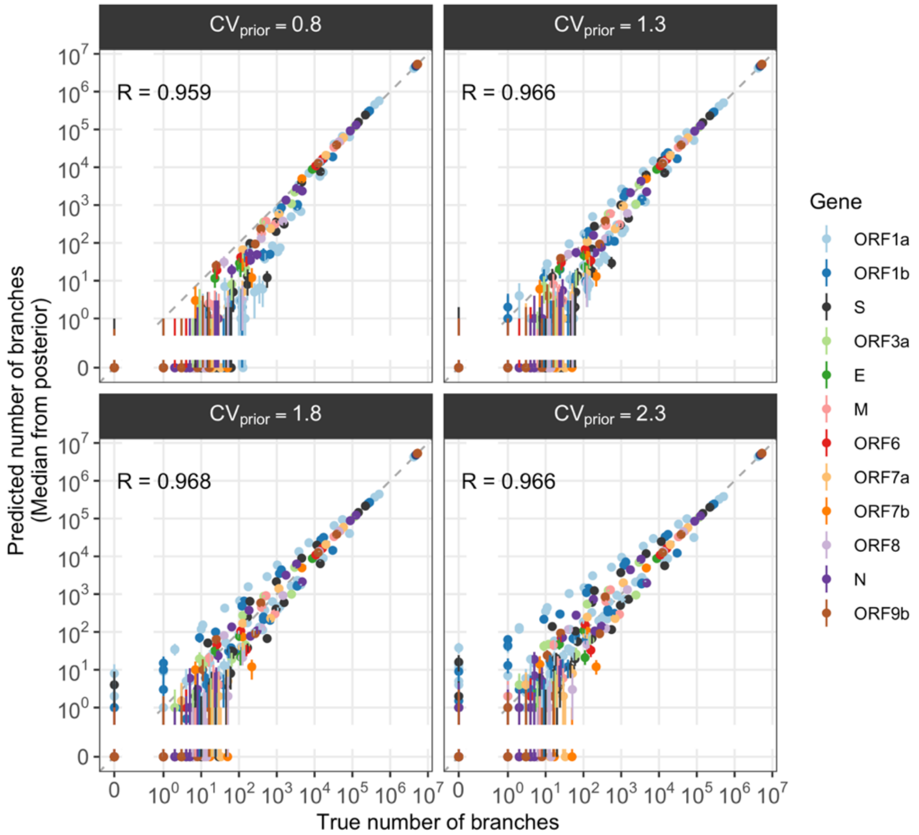


**Figure S14: Posterior predictive checks.** Comparison between the number of branches with a given number of nonsynonymous and synonymous mutations by gene across prior parametrizations. Points and vertical segments depict median and 95% credible intervals from posterior draws. For each prior parametrization (measured by the value of the coefficient of variation $CV_{prior}$), we also report the Pearson correlation coefficient $R$ between the log-transform predicted and observed number of branches, with a pseudo-count of 1 added prior to transformation (see Methods).

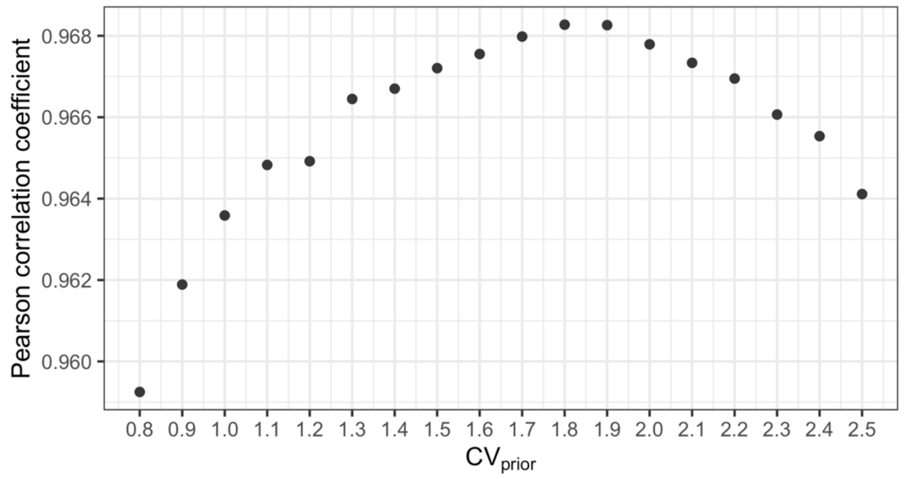


**Figure S15: Impact of prior parametrization on model performance.** Pearson correlation coefficient between the log-transformed predicted number of branches (median from the posterior) and the true number of branches with the same number of synonymous and nonsynonymous mutations across genes as a function of the coefficient of variation $CV_{prior}$ used on the prior parametrization. Prior to log-transformation, we add 1 to branch counts (see Methods).

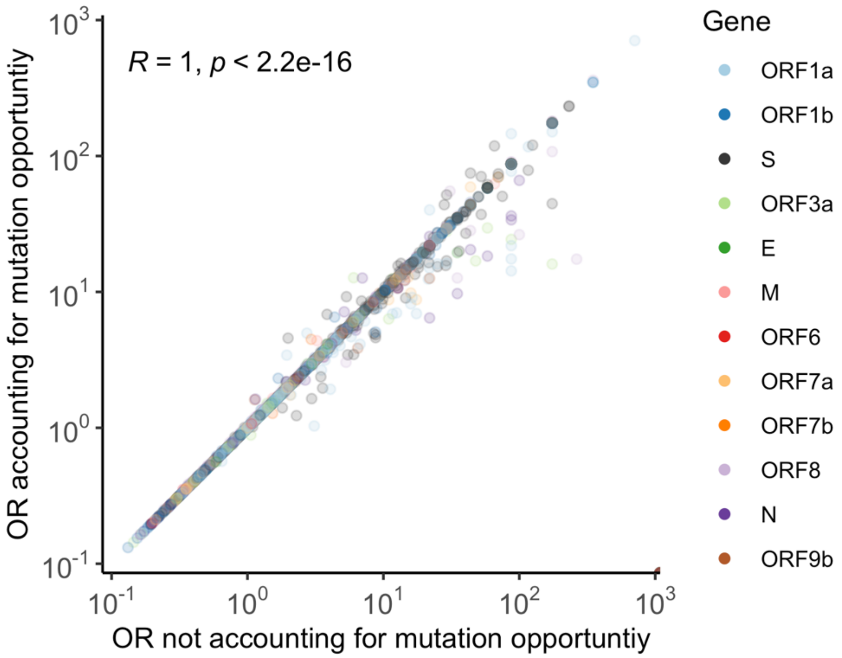


**Figure S16: Impact of accounting for mutation opportunity on the odds ratio for a mutation occurring in saltational vs non-saltational branches.** *R* corresponds to the Pearson correlation coefficient between the log ORs.

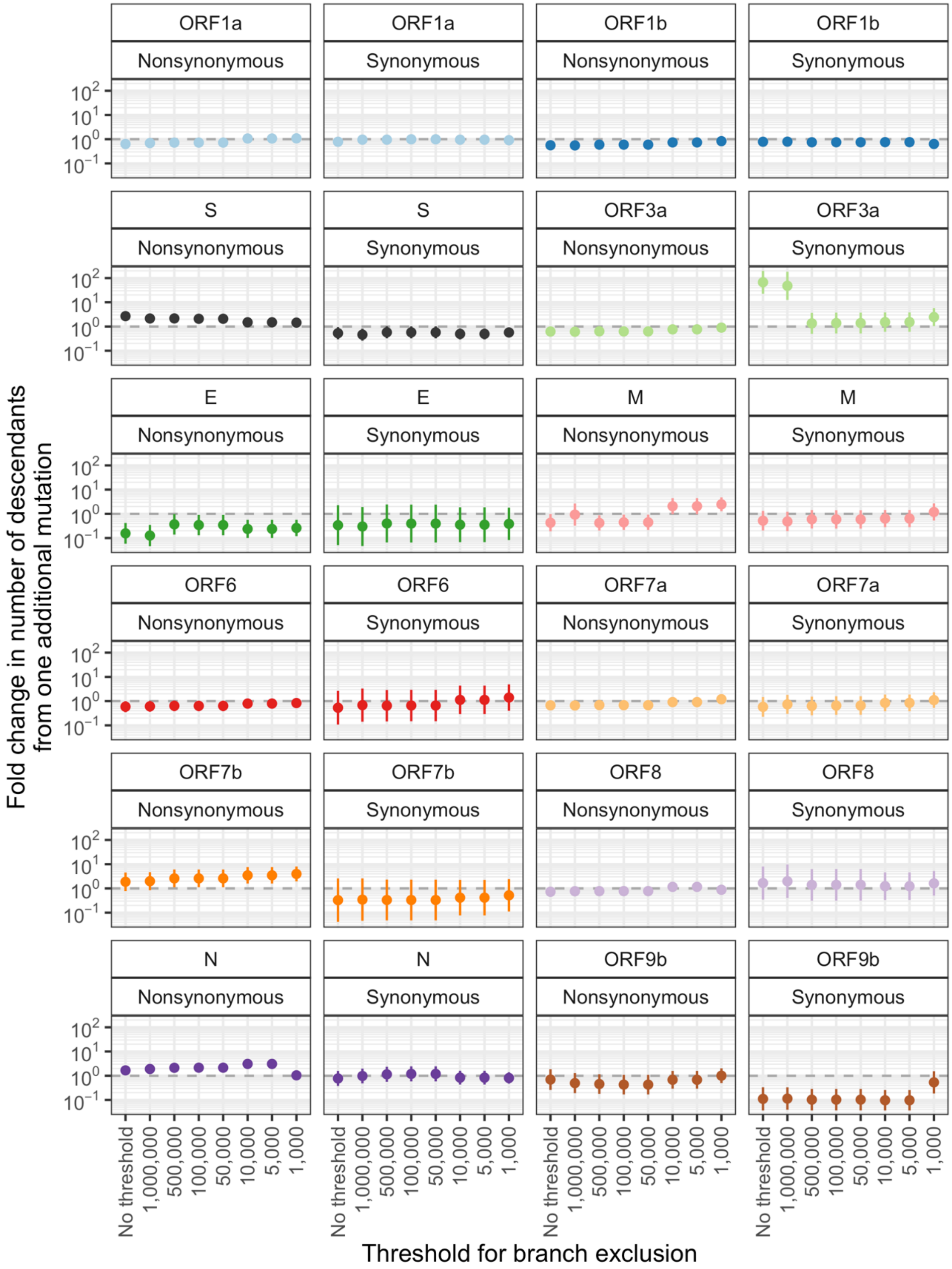


**Figure S17: Impact of branches with a high number of descendants on parameter estimates from a negative binomial regression.** We depict expected fold-change in the number of descendants from saltational branches from one additional mutation (by gene and mutation type) restricting the saltational branches included in the regression to those lying below the threshold depicted on the x-axis. Vertical segments indicate 95% confidence intervals.

**Table S1: List of positions that are more frequently mutated in saltational than non-saltational branches.** OR were computed using Fisher tests and p-values were adjusted for multiple testing (Benjamini-Hochberg corrections). We use a significance threshold of 0.05 for the adjusted p-value.

| Gene | Position | OR of position being mutated in saltational vs non-saltational branches (95% CI) | Saltational branches count with nonsynonymous mutations at position |
|---|---|---|---|
| S | 484 | 6.2 (4.6-8.2) | 51 |
| ORF8 | 121 | 4.1 (2.9-5.5) | 43 |
| S | 346 | 4.6 (3.3-6.3) | 41 |
| ORF8 | 120 | 6 (4.3-8.3) | 39 |
| S | 478 | 6.4 (4.4-9) | 35 |
| S | 446 | 2.7 (1.9-3.8) | 34 |
| S | 452 | 9.3 (6.4-13.2) | 34 |
| ORF1a | 1638 | 11.6 (7.9-16.5) | 33 |
| S | 681 | 7.5 (5.1-10.6) | 33 |
| S | 493 | 14.4 (9.4-21.1) | 28 |
| S | 936 | 3.1 (2-4.6) | 24 |
| ORF8 | 118 | 6.8 (4.3-10.3) | 23 |
| S | 444 | 3.2 (2-4.8) | 23 |
| S | 501 | 15.7 (9.8-24.2) | 23 |
| S | 9 | 5.1 (3.2-7.9) | 21 |
| S | 95 | 5.1 (3.2-7.9) | 21 |
| S | 494 | 2.7 (1.6-4.1) | 20 |
| S | 157 | 2.9 (1.7-4.5) | 19 |
| S | 440 | 25 (14.6-40.7) | 19 |
| S | 486 | 9.4 (5.6-14.9) | 19 |
| ORF1a | 1322 | 7.8 (4.6-12.5) | 18 |
| S | 152 | 3.9 (2.3-6.3) | 18 |
| S | 339 | 8.2 (4.8-13.2) | 18 |
| N | 203 | 3.6 (2.1-5.9) | 17 |
| ORF1a | 3255 | 13.5 (7.7-22.2) | 17 |
| ORF1a | 4398 | 2.3 (1.3-3.7) | 17 |
| ORF1a | 102 | 15.4 (8.6-25.7) | 16 |
| S | 257 | 4.7 (2.7-7.8) | 16 |
| S | 376 | 7.5 (4.3-12.4) | 16 |
| ORF1a | 1795 | 29.5 (15.9-51.4) | 15 |
| S | 340 | 8.3 (4.6-13.8) | 15 |
| S | 417 | 6.6 (3.7-11.1) | 15 |
| S | 490 | 4 (2.2-6.6) | 15 |
| M | 3 | 3 (1.7-5.1) | 14 |
| N | 204 | 2.9 (1.6-4.9) | 14 |
| ORF1a | 3209 | 3.3 (1.8-5.6) | 14 |
| S | 655 | 3 (1.6-5) | 14 |
| N | 234 | 4 (2.1-6.9) | 13 |
| ORF1a | 1323 | 10.4 (5.5-18.2) | 13 |
| ORF1b | 662 | 2.8 (1.5-4.8) | 13 |
| S | 210 | 9.2 (4.8-16) | 13 |

| | | | |
|---|---|---|---|
| S | 213 | 4.2 (2.2-7.2) | 13 |
| S | 214 | 2.8 (1.5-4.8) | 13 |
| S | 403 | 6.8 (3.6-11.8) | 13 |
| S | 477 | 3.1 (1.6-5.3) | 13 |
| ORF1a | 2980 | 5.3 (2.7-9.3) | 12 |
| ORF1a | 3201 | 5.3 (2.7-9.4) | 12 |
| S | 356 | 5.6 (2.9-9.9) | 12 |
| S | 252 | 3.6 (1.8-6.4) | 11 |
| S | 445 | 3.3 (1.6-5.9) | 11 |
| S | 450 | 8.3 (4.1-15.2) | 11 |
| ORF8 | 117 | 11.6 (5.4-21.9) | 10 |
| S | 64 | 4.8 (2.3-8.9) | 10 |
| S | 83 | 3.3 (1.6-6.1) | 10 |
| S | 460 | 5.1 (2.4-9.6) | 10 |
| ORF1a | 1500 | 3.5 (1.6-6.7) | 9 |
| ORF1a | 1639 | 8.6 (3.9-16.7) | 9 |
| ORF1a | 3653 | 10.7 (4.8-20.9) | 9 |
| ORF1a | 3915 | 3.4 (1.6-6.6) | 9 |
| ORF1a | 4097 | 6.4 (2.9-12.3) | 9 |
| ORF1a | 4165 | 8.3 (3.7-16.1) | 9 |
| S | 375 | 8.6 (3.9-16.6) | 9 |
| S | 498 | 25.8 (11.3-52.3) | 9 |
| S | 954 | 8.4 (3.8-16.3) | 9 |
| S | 449 | 18.9 (7.9-39.3) | 8 |
| E | 30 | 5 (2-10.5) | 7 |
| N | 235 | 6.8 (2.7-14.4) | 7 |
| ORF1a | 2771 | 7.3 (2.9-15.3) | 7 |
| ORF1b | 1000 | 5 (2-10.5) | 7 |
| ORF8 | 116 | 7.3 (2.9-15.5) | 7 |
| S | 373 | 5.8 (2.3-12.2) | 7 |
| S | 385 | 4.6 (1.8-9.7) | 7 |
| S | 481 | 4 (1.6-8.4) | 7 |
| S | 496 | 7 (2.8-14.8) | 7 |
| S | 1118 | 4.3 (1.7-8.9) | 7 |
| ORF1a | 1714 | 5.1 (1.8-11.3) | 6 |
| ORF1a | 2158 | 5.5 (2-12.3) | 6 |
| ORF1a | 2534 | 5.8 (2.1-12.8) | 6 |
| ORF9b | 93 | 6.8 (2.5-15.2) | 6 |
| S | 164 | 6 (2.2-13.3) | 6 |
| S | 368 | 9 (3.2-20.1) | 6 |
| S | 505 | 11.3 (4-25.5) | 6 |
| ORF1a | 47 | 6.4 (2.1-15.4) | 5 |
| ORF1a | 4167 | 12.5 (3.9-30.6) | 5 |
| S | 500 | 9.5 (3-23) | 5 |
| S | 764 | 18.2 (5.7-45.6) | 5 |
| S | 981 | 11.8 (3.7-28.9) | 5 |
| M | 19 | 8.4 (2.2-22.4) | 4 |
| ORF1a | 3651 | 58.3 (13.7-192.6) | 4 |
| ORF1a | 4324 | 14 (3.7-38.1) | 4 |

| | | | |
|---|---|---|---|
| ORF1b | 799 | 36.9 (9.1-110.7) | 4 |
| ORF1b | 800 | 21.9 (5.6-61.7) | 4 |
| S | 136 | 12.3 (3.2-33.2) | 4 |
| S | 568 | 10 (2.7-26.8) | 4 |
| S | 1190 | 12.7 (3.3-34.5) | 4 |
| N | 42 | 105 (16.3-545.1) | 3 |
| ORF1a | 1372 | 12.5 (2.5-39.1) | 3 |
| ORF1a | 2390 | 13.8 (2.7-43.5) | 3 |
| ORF1b | 804 | 29.2 (5.5-99.8) | 3 |
| S | 332 | 18.1 (3.5-58.4) | 3 |
| S | 1002 | 26.3 (5-88.6) | 3 |
| ORF1a | 2426 | 43.7 (4.5-217.9) | 2 |
| ORF1a | 3216 | 35 (3.7-163.4) | 2 |

**Table S2: Enrichment of nonsynonymous mutations at surface-accessible residues among mutations occurring on saltational branches.** We report odds ratios (OR) obtained using exact Fisher tests comparing the enrichment of Spike surface-accessible residues in saltational mutation across all mutations, mutations occurring at Spike residues and mutations occurring in the receptor-binding domain (RBD) within Spike.

| | **OR of a nonsynonymous mutation occurring at a Spike surface-accessible residue in saltational branches [95% CI]** |
|---|---|
| Across all mutations | 1.82 [1.72-1.92] |
| Across Spike mutations | 1.57 [1.43-1.73] |
| Across RBD mutations | 2.25 [1.77-2.89] |

**Table S3: List of nonsynonymous AA mutations that are more frequent in saltational than non-saltational branches.** OR were computed using Fisher tests and p-values were adjusted for multiple testing (Benjamini-Hochberg corrections). We use a significance threshold of 0.05 for the adjusted p-value. This table only contains mutations occurring at least 5 times on saltational branches.

| **Gene** | **Mutation** | **OR of mutation occurring in saltational vs non-saltational branches (95% CI)** | **Saltational branches count with mutation** |
|---|---|---|---|
| ORF8 | 121L | 4.86 (3.5-6.59) | 43 |
| ORF1a | 3829F | 1.86 (1.25-2.66) | 30 |
| ORF1a | 1638I | 13.51 (8.7-20.17) | 26 |
| ORF8 | 120L | 9.02 (5.72-13.59) | 24 |
| S | 484K | 15.02 (9.36-23.01) | 23 |
| S | 493Q | 43.79 (25.78-70.7) | 20 |
| S | 9L | 9.99 (6.03-15.67) | 20 |
| S | 452R | 13.47 (7.81-21.89) | 18 |
| S | 494P | 3.47 (2.05-5.52) | 18 |
| S | 95I | 3.46 (2.03-5.54) | 18 |
| ORF8 | 118V | 5.91 (3.41-9.56) | 17 |
| ORF1a | 4398L | 2.45 (1.42-3.93) | 17 |

| | | | |
|---|---|---|---|
| S | 501Y | 36.05 (19.97-61.48) | 17 |
| ORF1a | 3255I | 12.42 (6.89-20.96) | 16 |
| S | 681H | 9.84 (5.54-16.3) | 16 |
| ORF8 | 120V | 4.7 (2.61-7.81) | 15 |
| S | 346T | 4.56 (2.54-7.59) | 15 |
| ORF1a | 1795Q | 116.99 (55.13-241.07) | 14 |
| S | 446G | 13.4 (7.17-23.16) | 14 |
| S | 936H | 6.08 (3.29-10.32) | 14 |
| ORF1a | 1322I | 7.27 (3.83-12.62) | 13 |
| ORF1a | 102K | 36.4 (17.78-68.4) | 12 |
| S | 157L | 3.62 (1.86-6.37) | 12 |
| ORF1a | 3209V | 3.44 (1.77-6.06) | 12 |
| S | 440R | 87.49 (39.94-183.03) | 12 |
| S | 655Y | 2.88 (1.48-5.06) | 12 |
| ORF1b | 662S | 3.55 (1.82-6.25) | 12 |
| S | 478R | 4.13 (2.05-7.46) | 11 |
| S | 547K | 5 (2.47-9.05) | 11 |
| S | 152R | 6.33 (3-11.81) | 10 |
| S | 450D | 9.17 (4.32-17.23) | 10 |
| S | 477N | 11.61 (5.45-21.97) | 10 |
| S | 478K | 15.64 (7.14-30.79) | 10 |
| S | 484E | 30.69 (13.95-60.83) | 10 |
| S | 681R | 12.15 (5.68-23.12) | 10 |
| M | 82T | 12.24 (5.72-23.3) | 10 |
| ORF1a | 1639N | 11.51 (5.15-22.51) | 9 |
| N | 234I | 3.78 (1.72-7.25) | 9 |
| S | 346R | 7.46 (3.35-14.5) | 9 |
| S | 356T | 10.4 (4.66-20.27) | 9 |
| S | 376T | 16.11 (7.12-32.07) | 9 |
| N | 203K | 15.61 (6.54-32.12) | 8 |
| S | 257S | 6.67 (2.84-13.4) | 8 |
| ORF1a | 3653F | 28.63 (11.7-60.94) | 8 |
| S | 403K | 21.87 (9.06-45.75) | 8 |
| S | 460K | 12.22 (5.16-24.87) | 8 |
| S | 486F | 16.41 (6.79-34.33) | 8 |
| ORF1a | 1323G | 14.41 (5.62-31.02) | 7 |
| ORF1a | 1500Y | 3.42 (1.37-7.12) | 7 |
| S | 213E | 8.55 (3.38-18.05) | 7 |
| E | 30I | 6.01 (2.39-12.61) | 7 |
| S | 339G | 21.2 (8.18-46.44) | 7 |
| S | 452M | 7.77 (3.07-16.4) | 7 |
| S | 478T | 16.22 (6.34-34.81) | 7 |
| S | 481K | 11.64 (4.57-24.83) | 7 |
| S | 490S | 5.16 (2.05-10.79) | 7 |
| S | 83V | 19.96 (7.45-45.86) | 7 |
| ORF9b | 10P | 9.82 (3.53-22.07) | 6 |
| N | 13P | 9.26 (3.33-20.78) | 6 |
| S | 153T | 7.08 (2.56-15.79) | 6 |
| ORF1a | 1638N | 9.54 (3.43-21.45) | 6 |

| | | | |
|---|---|---|---|
| S | 190S | 7.79 (2.81-17.4) | 6 |
| N | 204G | 7.63 (2.75-17.05) | 6 |
| S | 210V | 12.45 (4.44-28.24) | 6 |
| S | 245N | 16.06 (5.69-36.88) | 6 |
| S | 257D | 4.28 (1.56-9.46) | 6 |
| S | 340D | 14.61 (5.19-33.39) | 6 |
| S | 346K | 3.89 (1.41-8.58) | 6 |
| ORF1a | 3571V | 3.97 (1.44-8.76) | 6 |
| S | 368I | 14.82 (5.27-33.79) | 6 |
| S | 375S | 12.45 (4.45-28.2) | 6 |
| ORF1a | 4097K | 8.74 (3.15-19.6) | 6 |
| S | 417T | 8.45 (3.04-18.92) | 6 |
| S | 449H | 33.97 (11.58-82.6) | 6 |
| S | 498R | 38.99 (13.12-97.1) | 6 |
| S | 70F | 3.87 (1.41-8.54) | 6 |
| S | 859N | 13.32 (4.75-30.27) | 6 |
| S | 954Q | 73.95 (23.49-200.57) | 6 |
| ORF1b | 1000L | 7.59 (2.41-18.33) | 5 |
| ORF1a | 1074T | 4.58 (1.47-10.88) | 5 |
| ORF1b | 1315R | 22.08 (6.76-56.53) | 5 |
| S | 158S | 19.96 (6.17-50.26) | 5 |
| ORF1a | 165Y | 4.56 (1.46-10.83) | 5 |
| S | 186L | 5.1 (1.63-12.12) | 5 |
| ORF9b | 18H | 5.06 (1.62-12.05) | 5 |
| N | 204R | 6.27 (2-14.99) | 5 |
| ORF1a | 2158I | 5.83 (1.87-13.92) | 5 |
| N | 235F | 19.26 (5.96-48.46) | 5 |
| S | 252G | 4.95 (1.57-11.96) | 5 |
| ORF1a | 2980N | 6.52 (2.08-15.6) | 5 |
| ORF1a | 3164H | 6.25 (2-14.94) | 5 |
| S | 385I | 5.34 (1.71-12.73) | 5 |
| ORF7a | 3F | 8.33 (2.65-20.07) | 5 |
| ORF1a | 4165A | 32.39 (9.74-85.55) | 5 |
| S | 417I | 35.1 (10.49-93.47) | 5 |
| S | 439K | 4.67 (1.5-11.1) | 5 |
| S | 440K | 42.36 (12.42-116.09) | 5 |
| S | 452Q | 5.82 (1.86-13.9) | 5 |
| S | 478E | 7.9 (2.51-18.99) | 5 |
| ORF7a | 49P | 8.24 (2.62-19.86) | 5 |
| ORF7a | 4L | 11.22 (3.54-27.32) | 5 |
| ORF1a | 542H | 4.73 (1.52-11.24) | 5 |
| ORF7a | 5V | 12.15 (3.83-29.69) | 5 |
| S | 64R | 5.76 (1.84-13.74) | 5 |
| S | 764K | 120.18 (30.15-445.25) | 5 |
| S | 899S | 4.84 (1.55-11.51) | 5 |

**Table S4: Association, among saltational branches, between having detected descendants and harboring a mutation enriched in saltational branches**

| Mutation | Number of occurrence in saltational branches with descendants | Number of occurrence in saltational branches without descendants | Odds ratio (95% CI) |
|---|---|---|---|
| S:452R | 12 | 6 | 4.8 (1.7-15.78) |
| ORF8:121L | 10 | 33 | 0.7 (0.3-1.49) |
| ORF1a:3255I | 9 | 7 | 3.1 (1-9.8) |
| ORF1a:3829F | 9 | 21 | 1 (0.4-2.33) |
| S:681H | 9 | 7 | 3.1 (1-9.8) |
| S:484K | 8 | 15 | 1.3 (0.5-3.21) |
| S:501Y | 8 | 9 | 2.1 (0.7-6.24) |
| M:82T | 8 | 2 | 9.6 (1.9-93.17) |
| S:936H | 8 | 6 | 3.2 (1-11.22) |
| S:95I | 8 | 10 | 1.9 (0.6-5.41) |
| S:477N | 7 | 3 | 5.6 (1.3-33.58) |
| S:9L | 7 | 13 | 1.3 (0.4-3.47) |
| ORF1a:102K | 6 | 6 | 2.4 (0.6-8.96) |
| ORF8:120L | 6 | 18 | 0.8 (0.3-2.08) |
| S:152R | 6 | 4 | 3.6 (0.8-17.33) |
| ORF1a:1795Q | 6 | 8 | 1.8 (0.5-5.9) |
| ORF1a:4398L | 6 | 11 | 1.3 (0.4-3.85) |
| S:450D | 6 | 4 | 3.6 (0.8-17.33) |
| S:478K | 6 | 4 | 3.6 (0.8-17.33) |
| S:478R | 6 | 5 | 2.9 (0.7-11.92) |
| S:493Q | 6 | 14 | 1 (0.3-2.83) |
| ORF1b:662S | 6 | 6 | 2.4 (0.6-8.96) |
| ORF1b:1000L | 5 | 0 | Inf (2.2-Inf) |
| S:157L | 5 | 7 | 1.7 (0.4-6.25) |
| ORF1a:1638I | 5 | 21 | 0.6 (0.2-1.54) |
| ORF1a:3209V | 5 | 7 | 1.7 (0.4-6.25) |
| S:356T | 5 | 4 | 3 (0.6-15.07) |
| ORF1a:3653F | 5 | 3 | 4 (0.8-25.69) |
| S:484E | 5 | 5 | 2.4 (0.5-10.4) |
| S:655Y | 5 | 7 | 1.7 (0.4-6.25) |
| ORF1a:1322I | 4 | 9 | 1.1 (0.2-3.79) |
| ORF1a:1638N | 4 | 2 | 4.8 (0.7-52.77) |
| S:213E | 4 | 3 | 3.2 (0.5-21.74) |
| N:234I | 4 | 5 | 1.9 (0.4-8.87) |
| S:346T | 4 | 11 | 0.9 (0.2-2.92) |
| S:440K | 4 | 1 | 9.5 (0.9-469.03) |
| S:440R | 4 | 8 | 1.2 (0.3-4.45) |
| S:446G | 4 | 10 | 0.9 (0.2-3.3) |
| S:460K | 4 | 4 | 2.4 (0.4-12.82) |
| S:490S | 4 | 3 | 3.2 (0.5-21.74) |
| S:547K | 4 | 7 | 1.4 (0.3-5.36) |
| S:899S | 4 | 1 | 9.5 (0.9-469.03) |
| ORF8:118V | 3 | 14 | 0.5 (0.1-1.82) |
| ORF8:120V | 3 | 12 | 0.6 (0.1-2.2) |
| ORF1a:1323G | 3 | 4 | 1.8 (0.3-10.56) |
| S:136F | 3 | 1 | 7.1 (0.6-374.31) |

| | | | |
|---|---|---|---|
| ORF1a:1500Y | 3 | 4 | 1.8 (0.3-10.56) |
| S:153T | 3 | 3 | 2.4 (0.3-17.78) |
| S:158S | 3 | 2 | 3.6 (0.4-42.79) |
| ORF1a:1714V | 3 | 1 | 7.1 (0.6-374.31) |
| S:190S | 3 | 3 | 2.4 (0.3-17.78) |
| N:203K | 3 | 5 | 1.4 (0.2-7.35) |
| N:204G | 3 | 3 | 2.4 (0.3-17.78) |
| S:213G | 3 | 1 | 7.1 (0.6-374.31) |
| N:213Y | 3 | 0 | Inf (1-Inf) |
| S:245N | 3 | 3 | 2.4 (0.3-17.78) |
| ORF1a:2534S | 3 | 1 | 7.1 (0.6-374.31) |
| ORF1b:2557R | 3 | 7 | 1 (0.2-4.47) |
| S:257S | 3 | 5 | 1.4 (0.2-7.35) |
| ORF1a:2980N | 3 | 2 | 3.6 (0.4-42.79) |
| S:346K | 3 | 3 | 2.4 (0.3-17.78) |
| S:356K | 3 | 0 | Inf (1-Inf) |
| ORF1a:3651S | 3 | 1 | 7.1 (0.6-374.31) |
| S:368I | 3 | 3 | 2.4 (0.3-17.78) |
| S:439K | 3 | 2 | 3.6 (0.4-42.79) |
| S:452M | 3 | 4 | 1.8 (0.3-10.56) |
| S:452Q | 3 | 2 | 3.6 (0.4-42.79) |
| S:494P | 3 | 15 | 0.5 (0.1-1.67) |
| S:64R | 3 | 2 | 3.6 (0.4-42.79) |
| S:681R | 3 | 7 | 1 (0.2-4.47) |
| S:764K | 3 | 2 | 3.6 (0.4-42.79) |
| S:859N | 3 | 3 | 2.4 (0.3-17.78) |
| ORF8:116A | 2 | 0 | Inf (0.4-Inf) |
| ORF8:117L | 2 | 1 | 4.7 (0.2-279.93) |
| ORF1a:1323A | 2 | 1 | 4.7 (0.2-279.93) |
| ORF1a:135R | 2 | 2 | 2.4 (0.2-32.8) |
| ORF1a:1639N | 2 | 7 | 0.7 (0.1-3.56) |
| S:164K | 2 | 1 | 4.7 (0.2-279.93) |
| S:183E | 2 | 1 | 4.7 (0.2-279.93) |
| S:186L | 2 | 3 | 1.6 (0.1-13.83) |
| ORF9b:18H | 2 | 3 | 1.6 (0.1-13.83) |
| N:203M | 2 | 1 | 4.7 (0.2-279.93) |
| S:210V | 2 | 4 | 1.2 (0.1-8.29) |
| ORF1a:2115I | 2 | 1 | 4.7 (0.2-279.93) |
| ORF1a:2140D | 2 | 1 | 4.7 (0.2-279.93) |
| ORF1a:2158I | 2 | 3 | 1.6 (0.1-13.83) |
| ORF1a:2364F | 2 | 2 | 2.4 (0.2-32.8) |
| S:257D | 2 | 4 | 1.2 (0.1-8.29) |
| S:262T | 2 | 1 | 4.7 (0.2-279.93) |
| ORF1a:2771R | 2 | 1 | 4.7 (0.2-279.93) |
| ORF1a:3164H | 2 | 3 | 1.6 (0.1-13.83) |
| S:332I | 2 | 0 | Inf (0.4-Inf) |
| S:339H | 2 | 0 | Inf (0.4-Inf) |
| S:346R | 2 | 7 | 0.7 (0.1-3.56) |
| ORF1a:3718A | 2 | 1 | 4.7 (0.2-279.93) |
| S:373P | 2 | 2 | 2.4 (0.2-32.8) |
| S:376A | 2 | 1 | 4.7 (0.2-279.93) |
| S:403K | 2 | 6 | 0.8 (0.1-4.43) |
| S:408S | 2 | 1 | 4.7 (0.2-279.93) |

| | | | |
|---|---|---|---|
| ORF1a:4097K | 2 | 4 | 1.2 (0.1-8.29) |
| ORF1a:4117N | 2 | 2 | 2.4 (0.2-32.8) |
| ORF1a:4161A | 2 | 1 | 4.7 (0.2-279.93) |
| S:417T | 2 | 4 | 1.2 (0.1-8.29) |
| S:449H | 2 | 4 | 1.2 (0.1-8.29) |
| S:478T | 2 | 5 | 0.9 (0.1-5.81) |
| S:484R | 2 | 2 | 2.4 (0.2-32.8) |
| S:486F | 2 | 6 | 0.8 (0.1-4.43) |
| S:486P | 2 | 1 | 4.7 (0.2-279.93) |
| S:493R | 2 | 1 | 4.7 (0.2-279.93) |
| S:498R | 2 | 4 | 1.2 (0.1-8.29) |
| ORF7a:49P | 2 | 3 | 1.6 (0.1-13.83) |
| E:51F | 2 | 1 | 4.7 (0.2-279.93) |
| ORF8:52R | 2 | 2 | 2.4 (0.2-32.8) |
| ORF9b:60A | 2 | 2 | 2.4 (0.2-32.8) |
| N:63G | 2 | 2 | 2.4 (0.2-32.8) |
| M:63T | 2 | 1 | 4.7 (0.2-279.93) |
| S:679K | 2 | 1 | 4.7 (0.2-279.93) |
| M:82S | 2 | 1 | 4.7 (0.2-279.93) |
| S:83V | 2 | 5 | 0.9 (0.1-5.81) |
| S:954H | 2 | 1 | 4.7 (0.2-279.93) |
| S:957H | 2 | 0 | Inf (0.4-Inf) |
| S:969K | 2 | 0 | Inf (0.4-Inf) |
| S:96Q | 2 | 1 | 4.7 (0.2-279.93) |
| S:981F | 2 | 0 | Inf (0.4-Inf) |
| S:1003I | 1 | 2 | 1.2 (0-22.79) |
| ORF1a:102V | 1 | 2 | 1.2 (0-22.79) |
| ORF7a:102V | 1 | 1 | 2.4 (0-185.87) |
| ORF1a:1074T | 1 | 4 | 0.6 (0-5.99) |
| ORF9b:10P | 1 | 5 | 0.5 (0-4.24) |
| S:1118D | 1 | 2 | 1.2 (0-22.79) |
| ORF1a:1140L | 1 | 2 | 1.2 (0-22.79) |
| ORF8:117F | 1 | 2 | 1.2 (0-22.79) |
| ORF8:118F | 1 | 1 | 2.4 (0-185.87) |
| ORF1a:1273G | 1 | 3 | 0.8 (0-9.85) |
| ORF1b:1315R | 1 | 4 | 0.6 (0-5.99) |
| ORF1a:1372A | 1 | 1 | 2.4 (0-185.87) |
| N:13P | 1 | 5 | 0.5 (0-4.24) |
| ORF1a:141A | 1 | 2 | 1.2 (0-22.79) |
| ORF1a:142P | 1 | 3 | 0.8 (0-9.85) |
| ORF1a:1436K | 1 | 1 | 2.4 (0-185.87) |
| S:152W | 1 | 2 | 1.2 (0-22.79) |
| ORF1a:1568T | 1 | 2 | 1.2 (0-22.79) |
| S:157F | 1 | 2 | 1.2 (0-22.79) |
| ORF1a:165Y | 1 | 4 | 0.6 (0-5.99) |
| S:188T | 1 | 1 | 2.4 (0-185.87) |
| ORF1a:1903Y | 1 | 3 | 0.8 (0-9.85) |
| N:199P | 1 | 1 | 2.4 (0-185.87) |
| M:19E | 1 | 2 | 1.2 (0-22.79) |
| N:202R | 1 | 1 | 2.4 (0-185.87) |
| N:205T | 1 | 2 | 1.2 (0-22.79) |
| S:210I | 1 | 2 | 1.2 (0-22.79) |
| S:210T | 1 | 3 | 0.8 (0-9.85) |

| | | | |
|---|---|---|---|
| ORF1a:2138A | 1 | 1 | 2.4 (0-185.87) |
| ORF1a:219Q | 1 | 1 | 2.4 (0-185.87) |
| N:234M | 1 | 3 | 0.8 (0-9.85) |
| N:235F | 1 | 4 | 0.6 (0-5.99) |
| S:252G | 1 | 4 | 0.6 (0-5.99) |
| ORF8:27Q | 1 | 2 | 1.2 (0-22.79) |
| ORF1a:2838L | 1 | 1 | 2.4 (0-185.87) |
| ORF7a:28I | 1 | 3 | 0.8 (0-9.85) |
| ORF1a:3090T | 1 | 3 | 0.8 (0-9.85) |
| ORF1a:3163S | 1 | 1 | 2.4 (0-185.87) |
| ORF1a:3201P | 1 | 2 | 1.2 (0-22.79) |
| S:339G | 1 | 6 | 0.4 (0-3.26) |
| S:339Y | 1 | 3 | 0.8 (0-9.85) |
| S:340D | 1 | 5 | 0.5 (0-4.24) |
| ORF1a:3573E | 1 | 1 | 2.4 (0-185.87) |
| ORF1a:3580Q | 1 | 1 | 2.4 (0-185.87) |
| S:385I | 1 | 4 | 0.6 (0-5.99) |
| ORF1a:3949N | 1 | 3 | 0.8 (0-9.85) |
| ORF7a:3F | 1 | 4 | 0.6 (0-5.99) |
| ORF1a:4167S | 1 | 2 | 1.2 (0-22.79) |
| S:445H | 1 | 1 | 2.4 (0-185.87) |
| S:445P | 1 | 1 | 2.4 (0-185.87) |
| S:446R | 1 | 2 | 1.2 (0-22.79) |
| S:449N | 1 | 1 | 2.4 (0-185.87) |
| S:452L | 1 | 2 | 1.2 (0-22.79) |
| ORF1a:47K | 1 | 3 | 0.8 (0-9.85) |
| S:481K | 1 | 6 | 0.4 (0-3.26) |
| S:496S | 1 | 3 | 0.8 (0-9.85) |
| ORF7a:4L | 1 | 4 | 0.6 (0-5.99) |
| S:501T | 1 | 3 | 0.8 (0-9.85) |
| S:505H | 1 | 2 | 1.2 (0-22.79) |
| ORF1a:542H | 1 | 4 | 0.6 (0-5.99) |
| S:568N | 1 | 2 | 1.2 (0-22.79) |
| ORF3a:57Q | 1 | 2 | 1.2 (0-22.79) |
| ORF7a:5V | 1 | 4 | 0.6 (0-5.99) |
| S:67A | 1 | 3 | 0.8 (0-9.85) |
| S:716T | 1 | 2 | 1.2 (0-22.79) |
| ORF8:73Y | 1 | 2 | 1.2 (0-22.79) |
| ORF1b:797A | 1 | 3 | 0.8 (0-9.85) |
| ORF1b:798Q | 1 | 1 | 2.4 (0-185.87) |
| ORF1b:799R | 1 | 2 | 1.2 (0-22.79) |
| ORF1b:800R | 1 | 2 | 1.2 (0-22.79) |
| ORF1b:803I | 1 | 1 | 2.4 (0-185.87) |
| ORF1b:804G | 1 | 2 | 1.2 (0-22.79) |
| ORF7a:89K | 1 | 2 | 1.2 (0-22.79) |
| S:954Q | 1 | 5 | 0.5 (0-4.24) |
| S:976A | 1 | 1 | 2.4 (0-185.87) |
| ORF7a:97C | 1 | 2 | 1.2 (0-22.79) |
| ORF1b:991T | 1 | 1 | 2.4 (0-185.87) |
| S:1002H | 0 | 2 | 0 (0-12.61) |
| ORF3a:10L | 0 | 2 | 0 (0-12.61) |
| S:1187K | 0 | 2 | 0 (0-12.61) |
| S:1188V | 0 | 2 | 0 (0-12.61) |

| | | | |
|---|---|---|---|
| ORF8:118R | 0 | 2 | 0 (0-12.61) |
| ORF8:119S | 0 | 3 | 0 (0-5.73) |
| ORF8:11T | 0 | 2 | 0 (0-12.61) |
| ORF1a:1322A | 0 | 4 | 0 (0-3.58) |
| N:13R | 0 | 2 | 0 (0-12.61) |
| ORF1a:154H | 0 | 2 | 0 (0-12.61) |
| ORF1a:1709H | 0 | 2 | 0 (0-12.61) |
| S:187T | 0 | 2 | 0 (0-12.61) |
| N:204R | 0 | 5 | 0 (0-2.58) |
| ORF1b:2314V | 0 | 2 | 0 (0-12.61) |
| ORF1a:2390S | 0 | 2 | 0 (0-12.61) |
| ORF1a:2430A | 0 | 3 | 0 (0-5.73) |
| ORF1b:2653D | 0 | 2 | 0 (0-12.61) |
| E:30I | 0 | 7 | 0 (0-1.64) |
| ORF1a:3161I | 0 | 2 | 0 (0-12.61) |
| ORF1a:3312L | 0 | 2 | 0 (0-12.61) |
| ORF1a:3350I | 0 | 2 | 0 (0-12.61) |
| S:339N | 0 | 2 | 0 (0-12.61) |
| ORF1a:3571V | 0 | 6 | 0 (0-2.01) |
| S:357K | 0 | 4 | 0 (0-3.58) |
| ORF1a:3606V | 0 | 2 | 0 (0-12.61) |
| S:372T | 0 | 2 | 0 (0-12.61) |
| S:373S | 0 | 3 | 0 (0-5.73) |
| S:375S | 0 | 6 | 0 (0-2.01) |
| S:376T | 0 | 9 | 0 (0-1.19) |
| ORF1a:3915I | 0 | 3 | 0 (0-5.73) |
| S:403S | 0 | 2 | 0 (0-12.61) |
| S:405D | 0 | 3 | 0 (0-5.73) |
| ORF1a:4165A | 0 | 5 | 0 (0-2.58) |
| S:417I | 0 | 5 | 0 (0-2.58) |
| ORF1a:4206H | 0 | 3 | 0 (0-5.73) |
| ORF1a:4223S | 0 | 4 | 0 (0-3.58) |
| ORF1a:4324T | 0 | 3 | 0 (0-5.73) |
| ORF1a:4358K | 0 | 4 | 0 (0-3.58) |
| S:453F | 0 | 3 | 0 (0-5.73) |
| S:478E | 0 | 5 | 0 (0-2.58) |
| S:500N | 0 | 2 | 0 (0-12.61) |
| S:500S | 0 | 3 | 0 (0-5.73) |
| S:532K | 0 | 2 | 0 (0-12.61) |
| ORF1a:549Y | 0 | 2 | 0 (0-12.61) |
| S:570A | 0 | 3 | 0 (0-5.73) |
| ORF1a:571G | 0 | 4 | 0 (0-3.58) |
| ORF7a:60D | 0 | 2 | 0 (0-12.61) |
| S:65S | 0 | 2 | 0 (0-12.61) |
| S:681P | 0 | 3 | 0 (0-5.73) |
| S:68L | 0 | 3 | 0 (0-5.73) |
| S:69P | 0 | 4 | 0 (0-3.58) |
| E:6L | 0 | 2 | 0 (0-12.61) |
| S:70F | 0 | 6 | 0 (0-2.01) |
| S:73S | 0 | 4 | 0 (0-3.58) |
| ORF1a:74A | 0 | 2 | 0 (0-12.61) |
| ORF1a:827E | 0 | 2 | 0 (0-12.61) |
| ORF9b:93L | 0 | 4 | 0 (0-3.58) |

| | | | |
|---|---|---|---|
| S:95N | 0 | 3 | 0 (0-5.73) |
| N:96V | 0 | 4 | 0 (0-3.58) |
| S:67A | 1 | 3 | 0.8 (0-9.85) |
| S:716T | 1 | 2 | 1.2 (0-22.79) |
| ORF8:73Y | 1 | 2 | 1.2 (0-22.79) |
| ORF1b:797A | 1 | 3 | 0.8 (0-9.85) |
| ORF1b:798Q | 1 | 1 | 2.4 (0-185.87) |
| ORF1b:799R | 1 | 2 | 1.2 (0-22.79) |
| ORF1b:800R | 1 | 2 | 1.2 (0-22.79) |
| ORF1b:803I | 1 | 1 | 2.4 (0-185.87) |
| ORF1b:804G | 1 | 2 | 1.2 (0-22.79) |
| ORF7a:89K | 1 | 2 | 1.2 (0-22.79) |
| S:954Q | 1 | 5 | 0.5 (0-4.24) |
| S:976A | 1 | 1 | 2.4 (0-185.87) |
| ORF7a:97C | 1 | 2 | 1.2 (0-22.79) |
| ORF1b:991T | 1 | 1 | 2.4 (0-185.87) |
| S:1002H | 0 | 2 | 0 (0-12.61) |
| ORF3a:10L | 0 | 2 | 0 (0-12.61) |
| S:1187K | 0 | 2 | 0 (0-12.61) |
| S:1188V | 0 | 2 | 0 (0-12.61) |
| ORF8:118R | 0 | 2 | 0 (0-12.61) |
| ORF8:119S | 0 | 3 | 0 (0-5.73) |
| ORF8:11T | 0 | 2 | 0 (0-12.61) |
| ORF1a:1322A | 0 | 4 | 0 (0-3.58) |
| N:13R | 0 | 2 | 0 (0-12.61) |
| ORF1a:154H | 0 | 2 | 0 (0-12.61) |
| ORF1a:1709H | 0 | 2 | 0 (0-12.61) |
| S:187T | 0 | 2 | 0 (0-12.61) |
| N:204R | 0 | 5 | 0 (0-2.58) |
| ORF1b:2314V | 0 | 2 | 0 (0-12.61) |
| ORF1a:2390S | 0 | 2 | 0 (0-12.61) |
| ORF1a:2430A | 0 | 3 | 0 (0-5.73) |
| ORF1b:2653D | 0 | 2 | 0 (0-12.61) |
| E:30I | 0 | 7 | 0 (0-1.64) |
| ORF1a:3161I | 0 | 2 | 0 (0-12.61) |
| ORF1a:3312L | 0 | 2 | 0 (0-12.61) |
| ORF1a:3350I | 0 | 2 | 0 (0-12.61) |
| S:339N | 0 | 2 | 0 (0-12.61) |
| ORF1a:3571V | 0 | 6 | 0 (0-2.01) |
| S:357K | 0 | 4 | 0 (0-3.58) |
| ORF1a:3606V | 0 | 2 | 0 (0-12.61) |
| S:372T | 0 | 2 | 0 (0-12.61) |
| S:373S | 0 | 3 | 0 (0-5.73) |
| S:375S | 0 | 6 | 0 (0-2.01) |
| S:376T | 0 | 9 | 0 (0-1.19) |
| ORF1a:3915I | 0 | 3 | 0 (0-5.73) |
| S:403S | 0 | 2 | 0 (0-12.61) |
| S:405D | 0 | 3 | 0 (0-5.73) |
| ORF1a:4165A | 0 | 5 | 0 (0-2.58) |
| S:417I | 0 | 5 | 0 (0-2.58) |
| ORF1a:4206H | 0 | 3 | 0 (0-5.73) |
| ORF1a:4223S | 0 | 4 | 0 (0-3.58) |
| ORF1a:4324T | 0 | 3 | 0 (0-5.73) |

| | | | |
|---|---|---|---|
| ORF1a:4358K | 0 | 4 | 0 (0-3.58) |
| S:453F | 0 | 3 | 0 (0-5.73) |
| S:478E | 0 | 5 | 0 (0-2.58) |
| S:500N | 0 | 2 | 0 (0-12.61) |
| S:500S | 0 | 3 | 0 (0-5.73) |
| S:532K | 0 | 2 | 0 (0-12.61) |
| ORF1a:549Y | 0 | 2 | 0 (0-12.61) |
| S:570A | 0 | 3 | 0 (0-5.73) |
| ORF1a:571G | 0 | 4 | 0 (0-3.58) |
| ORF7a:60D | 0 | 2 | 0 (0-12.61) |
| S:65S | 0 | 2 | 0 (0-12.61) |
| S:681P | 0 | 3 | 0 (0-5.73) |
| S:68L | 0 | 3 | 0 (0-5.73) |
| S:69P | 0 | 4 | 0 (0-3.58) |
| E:6L | 0 | 2 | 0 (0-12.61) |
| S:70F | 0 | 6 | 0 (0-2.01) |
| S:73S | 0 | 4 | 0 (0-3.58) |
| ORF1a:74A | 0 | 2 | 0 (0-12.61) |
| ORF1a:827E | 0 | 2 | 0 (0-12.61) |
| ORF9b:93L | 0 | 4 | 0 (0-3.58) |
| S:95N | 0 | 3 | 0 (0-5.73) |
| N:96V | 0 | 4 | 0 (0-3.58) |

**Table S5: Summary of recurrent amino-acid mutations found in persistent SARS-CoV-2 infections.** Summary of recurrent nonsynonymous mutations reported in chronic SARS-CoV-2 infections[1–4] or suspected chronic SARS-CoV-2 infections[5]. We report mutations observed in at least 3 infections except in Harari et al.[4] (Nature Medicine) and Ghafari et al.[3] (Nature) where we report mutations observed in at least 2 persistent infections (due to the lower sample size and number of mutations satisfying this criterion).

| Mutation | Source |
|---|---|
| S:E340K | Khurana et al. medRxiv. 2025.[1]<br>Harari et al. Nature Communications. 2024.[5] |
| S:N405D | Khurana et al. medRxiv. 2025.[1] |
| ORF1a:L3829F | Khurana et al. medRxiv. 2025.[1] |
| ORF1b:G662S | Khurana et al. medRxiv. 2025.[1] |
| ORF1a:T1638I | Khurana et al. medRxiv. 2025.[1]<br>Wilkinson et al. Virus Evolution. 2022.[2]<br>Ghafari et al. Nature. 2024.[3]<br>Harari et al. Nature Medicine. 2022.[4]<br>Harari et al. Nature Communications. 2024.[5] |
| S:R346T | Khurana et al. medRxiv. 2025.[1]<br>Harari et al. Nature Communications. 2024.[5] |
| S:K444R | Khurana et al. medRxiv. 2025.[1] |
| N:R93R | Khurana et al. medRxiv. 2025.[1] |
| E:T30I | Khurana et al. medRxiv. 2025.[1]<br>Wilkinson et al. Virus Evolution. 2022.[2]<br>Harari et al. Nature Medicine. 2022.[4]<br>Harari et al. Nature Communications. 2024.[5] |
| S:N417I | Khurana et al. medRxiv. 2025.[1] |
| ORF1a:D1639N | Khurana et al. medRxiv. 2025.[1] |
| S:L452M | Khurana et al. medRxiv. 2025.[1] |
| ORF1b:V783I | Khurana et al. medRxiv. 2025.[1] |
| S:R346I | Khurana et al. medRxiv. 2025.[1] |
| S:D405N | Khurana et al. medRxiv. 2025.[1] |
| S:K764N | Khurana et al. medRxiv. 2025.[1] |
| ORF1a:T4175I | Khurana et al. medRxiv. 2025.[1]<br>Ghafari et al. Nature. 2024.[3] |
| S:T19I | Khurana et al. medRxiv. 2025.[1] |
| ORF3a:S74S | Khurana et al. medRxiv. 2025.[1] |
| N:P326L | Khurana et al. medRxiv. 2025.[1] |
| ORF1a:T1322I | Khurana et al. medRxiv. 2025.[1]<br>Harari et al. Nature Medicine. 2022.[4] |
| S:E340Q | Khurana et al. medRxiv. 2025.[1]<br>Harari et al. Nature Communications. 2024.[5] |
| S:K444N | Khurana et al. medRxiv. 2025.[1]<br>Harari et al. Nature Communications. 2024.[5] |
| S:I19T | Khurana et al. medRxiv. 2025.[1] |
| S:S494P | Khurana et al. medRxiv. 2025.[1]<br>Harari et al. Nature Medicine. 2022.[4] |
| S:K440R | Khurana et al. medRxiv. 2025.[1] |
| ORF1a:S4398L | Khurana et al. medRxiv. 2025.[1] |
| S:P9L | Khurana et al. medRxiv. 2025.[1] |

| | |
|---|---|
| | Harari et al. Nature Medicine. 2022.[4] |
| S:T95I | Khurana et al. medRxiv. 2025.[1]<br>Wilkinson et al. Virus Evolution. 2022.[2]<br>Harari et al. Nature Communications. 2024.[5] |
| ORF1a:T1538I | Khurana et al. medRxiv. 2025.[1] |
| ORF1a:A2529V | Khurana et al. medRxiv. 2025.[1] |
| ORF1a:E102K | Khurana et al. medRxiv. 2025.[1] |
| ORF1a:G3072C | Khurana et al. medRxiv. 2025.[1] |
| S:E484K | Wilkinson et al. Virus Evolution. 2022.[2]<br>Harari et al. Nature Medicine. 2022.[4] |
| M:H125Y | Wilkinson et al. Virus Evolution. 2022.[2]<br>Harari et al. Nature Medicine. 2022.[4] |
| ORF1a:T3058I | Wilkinson et al. Virus Evolution. 2022.[2]<br>Harari et al. Nature Medicine. 2022.[4] |
| S:Q493K | Wilkinson et al. Virus Evolution. 2022.[2] |
| S:S13I | Wilkinson et al. Virus Evolution. 2022.[2]<br>Harari et al. Nature Medicine. 2022.[4] |
| ORF1b:V783I | Wilkinson et al. Virus Evolution. 2022.[2]<br>Harari et al. Nature Medicine. 2022.[4] |
| ORF1a:P1640L | Wilkinson et al. Virus Evolution. 2022.[2]<br>Harari et al. Nature Medicine. 2022.[4] |
| ORF1a:T1322P | Wilkinson et al. Virus Evolution. 2022.[2] |
| S:L452R | Wilkinson et al. Virus Evolution. 2022.[2]<br>Ghafari et al. Nature. 2024.[3] |
| S:Q498R | Wilkinson et al. Virus Evolution. 2022.[2] |
| S:E484G | Wilkinson et al. Virus Evolution. 2022.[2] |
| S:F486I | Wilkinson et al. Virus Evolution. 2022.[2] |
| S:G142V | Wilkinson et al. Virus Evolution. 2022.[2] |
| ORF1a:K1795Q | Wilkinson et al. Virus Evolution. 2022.[2]<br>Harari et al. Nature Communications. 2024.[5] |
| S:F490L | Wilkinson et al. Virus Evolution. 2022.[2] |
| ORF1a:T4175I | Ghafari et al. Nature. 2024.[3] |
| ORF1b:L1504F | Ghafari et al. Nature. 2024.[3] |
| S:T376A | Ghafari et al. Nature. 2024.[3] |
| S:T547K | Ghafari et al. Nature. 2024.[3] |
| S:R190K | Harari et al. Nature Medicine. 2022.[4] |
| S:N501Y | Harari et al. Nature Medicine. 2022.[4] |
| S:E484Q | Harari et al. Nature Medicine. 2022.[4] |
| ORF7a:A105V | Harari et al. Nature Medicine. 2022.[4] |
| ORF8:T11I | Harari et al. Nature Medicine. 2022.[4] |
| ORF1a:V1887I | Harari et al. Nature Communications. 2024.[5] |
| S:S371F | Harari et al. Nature Communications. 2024.[5] |
| S:A701V | Harari et al. Nature Communications. 2024.[5] |
| S:K356T | Harari et al. Nature Communications. 2024.[5] |
| S:E340D | Harari et al. Nature Communications. 2024.[5] |
| S:L455S | Harari et al. Nature Communications. 2024.[5] |
| S:N460K | Harari et al. Nature Communications. 2024.[5] |
| ORF1a:P1803S | Harari et al. Nature Communications. 2024.[5] |
| ORF3a:W131C | Harari et al. Nature Communications. 2024.[5] |

| | |
|---|---|
| S:E340A | Harari et al. Nature Communications. 2024.[5] |
| S:G446V | Harari et al. Nature Communications. 2024.[5] |
| S:P337S | Harari et al. Nature Communications. 2024.[5] |
| ORF3a :E239Q | Harari et al. Nature Communications. 2024.[5] |

**Table S6: Summary of recurrent amino-acid mutations reported in minks infected with SARS-CoV-2.** We report the 5 mutations found to be strong candidate for mink-specific adaptation by Tan et al.[6], 3 mutations found to recur in minks in Zhou et al.[7] For Iglesias-Caballero et al.[8], we extracted recurrent nonsynonymous mutations across clusters from their second figure.

| Mutation | Source |
|---|---|
| S:Y453F | Zhou et al. Cell Reports. 2022.[7]<br>Tan et al. Nature Communications. 2025.[6] |
| S:F486L | Zhou et al. Cell Reports. 2022.[7]<br>Tan et al. Nature Communications. 2025.[6]<br>Iglesias-Caballero et al. Int. J. Nol. Sci. 2024.[8] |
| S:N501T | Zhou et al. Cell Reports. 2022.[7]<br>Tan et al. Nature Communications. 2025.[6]<br>Iglesias-Caballero et al. Int. J. Nol. Sci. 2024.[8] |
| ORF1a:G1477E | Tan et al. Nature Communications. 2025.[6]<br>Iglesias-Caballero et al. Int. J. Nol. Sci. 2024.[8] |
| ORF3a:L219V | Tan et al. Nature Communications. 2025.[6] |
| ORF1a:T265I | Iglesias-Caballero et al. Int. J. Nol. Sci. 2024.[8] |

**Table S7: Summary of recurrent amino-acid mutations reported in deers infected with SARS-CoV-2.** We report a mutation found to be a strong candidate for deer-specific adaptation by Tan et al. [4], 2 mutations found to occur at least 3 times independently in Marques et al.[9] and mutations found in 3 independent samples in Feng et al.[10]

| Mutation | Source |
|---|---|
| ORF1a:L1853F | Tan et al. Nature Communications. 2025.[6] |
| S:V252G | Marques et al. PLoS Pathogens. 2025.[9] |
| ORF1a:S3149F | Marques et al. PLoS Pathogens. 2025.[9] |
| ORF1a:H110Y | Feng et al. Nature Communications. 2023.[10] |
| ORF1a:S443P | Feng et al. Nature Communications. 2023.[10] |
| ORF1a:S944L | Feng et al. Nature Communications. 2023.[10] |
| ORF1a:T1000I | Feng et al. Nature Communications. 2023.[10] |
| ORF1a:P1640S | Feng et al. Nature Communications. 2023.[10] |
| ORF1a:T1854I | Feng et al. Nature Communications. 2023.[10] |
| ORF1a:T1881I | Feng et al. Nature Communications. 2023.[10] |
| ORF1a:T2016I | Feng et al. Nature Communications. 2023.[10] |
| ORF1a:S2103F | Feng et al. Nature Communications. 2023.[10] |
| ORF1a:S2224F | Feng et al. Nature Communications. 2023.[10] |
| ORF1a:S2242F | Feng et al. Nature Communications. 2023.[10] |
| ORF1a:S2255F | Feng et al. Nature Communications. 2023.[10] |
| ORF1a:S2273F | Feng et al. Nature Communications. 2023.[10] |
| ORF1a:T2283I | Feng et al. Nature Communications. 2023.[10] |
| ORF1a:H2357Y | Feng et al. Nature Communications. 2023.[10] |
| ORF1a:K2511N | Feng et al. Nature Communications. 2023.[10] |

| | |
|---|---|
| ORF1a:V3017I | Feng et al. Nature Communications. 2023.[10] |
| ORF1a:H3076Y | Feng et al. Nature Communications. 2023.[10] |
| ORF1a:T3082I | Feng et al. Nature Communications. 2023.[10] |
| ORF1a:S3099L | Feng et al. Nature Communications. 2023.[10] |
| ORF1a:L3116F | Feng et al. Nature Communications. 2023.[10] |
| ORF1a:S3149F | Feng et al. Nature Communications. 2023.[10] |
| ORF1a:T3150I | Feng et al. Nature Communications. 2023.[10] |
| ORF1a:A3548V | Feng et al. Nature Communications. 2023.[10] |
| ORF1a:T3646I | Feng et al. Nature Communications. 2023.[10] |
| ORF1a:L3808F | Feng et al. Nature Communications. 2023.[10] |
| ORF1a:T3904I | Feng et al. Nature Communications. 2023.[10] |
| ORF1a:L4111F | Feng et al. Nature Communications. 2023.[10] |
| ORF1b:A88V | Feng et al. Nature Communications. 2023.[10] |
| ORF1b:S2338L | Feng et al. Nature Communications. 2023.[10] |
| ORF1b:S2339F | Feng et al. Nature Communications. 2023.[10] |
| ORF1b:T2376I | Feng et al. Nature Communications. 2023.[10] |
| S:T22I | Feng et al. Nature Communications. 2023.[10] |
| S:T29I | Feng et al. Nature Communications. 2023.[10] |
| S:H146Y | Feng et al. Nature Communications. 2023.[10] |
| S:H245Y | Feng et al. Nature Communications. 2023.[10] |
| S:S640F | Feng et al. Nature Communications. 2023.[10] |
| S:S680F | Feng et al. Nature Communications. 2023.[10] |
| S:L1203F | Feng et al. Nature Communications. 2023.[10] |
| ORF3a:A54V | Feng et al. Nature Communications. 2023.[10] |
| ORF3a: L108F | Feng et al. Nature Communications. 2023.[10] |
| ORF7a:T39I | Feng et al. Nature Communications. 2023.[10] |
| ORF7b:S5L | Feng et al. Nature Communications. 2023.[10] |
| ORF8:S43F | Feng et al. Nature Communications. 2023.[10] |
| ORF8:K68E | Feng et al. Nature Communications. 2023.[10] |
| N:T271I | Feng et al. Nature Communications. 2023.[10] |
| N:T334I | Feng et al. Nature Communications. 2023.[10] |

#### References


1. Khurana, M. P. *et al.* Large-scale genomic surveillance reveals immunosuppression drives mutation dynamics in persistent SARS-CoV-2 infections. *medRxiv* (2025) doi:10.1101/2025.02.10.25321987.
2. Wilkinson, S. A. J. *et al.* Recurrent SARS-CoV-2 mutations in immunodeficient patients. *Virus Evol.* **8**, veac050 (2022).
3. Ghafari, M. *et al.* Prevalence of persistent SARS-CoV-2 in a large community surveillance study. *Nature* **626**, 1094–1101 (2024).
4. Harari, S. *et al.* Drivers of adaptive evolution during chronic SARS-CoV-2 infections. *Nat. Med.* **28**, 1501–1508 (2022).
5. Harari, S., Miller, D., Fleishon, S., Burstein, D. & Stern, A. Using big sequencing data to identify chronic SARS-Coronavirus-2 infections. *Nat. Commun.* **15**, 648 (2024).
6. Tan, C. C. S. *et al.* Transmission of SARS-CoV-2 from humans to animals and potential host adaptation. *Nat. Commun.* **13**, 2988 (2022).
7. Zhou, J. *et al.* Mutations that adapt SARS-CoV-2 to mink or ferret do not increase fitness in the human airway. *Cell Rep.* **38**, 110344 (2022).
8. Iglesias-Caballero, M. *et al.* Genomic context of SARS-CoV-2 outbreaks in farmed mink in Spain during pandemic: Unveiling host adaptation mechanisms. *Int. J. Mol. Sci.* **25**, 5499 (2024).
9. Marques, A. D. *et al.* Evolution of SARS-CoV-2 in white-tailed deer in Pennsylvania 2021-2024. *PLoS Pathog.* **21**, e1012883 (2025).
10. Feng, A. *et al.* Transmission of SARS-CoV-2 in free-ranging white-tailed deer in the United States. *Nat. Commun.* **14**, 4078 (2023).
